# Optical interrogation of sympathetic neuronal effects on macroscopic cardiac monolayer dynamics

**DOI:** 10.1101/717991

**Authors:** R.A.B. Burton, J. Tomek, C.M. Ambrosi, H.E. Larsen, A.R. Sharkey, R.A. Capel, S. Bilton, A. Klimas, G. Stephens, D. Li, G. Gallone, N. Herring, E. Mann, A. Kumar, H. Kramer, E. Entcheva, D.J. Paterson, G. Bub

## Abstract

Alterations in autonomic function are known to occur in cardiac conditions including sudden cardiac death. Cardiac stimulation via sympathetic neurons can potentially trigger arrhythmias. Dissecting direct neural-cardiac interactions at the cellular level is technically challenging and understudied due to the lack of experimental model systems and methodologies. Here we demonstrate the utility of optical interrogation of sympathetic neurons and their effects on macroscopic cardiac monolayer dynamics to address research targets such as the effects of adrenergic stimulation via the release of neurotransmitters, the effect of neuronal numbers on cardiac wave behaviour and the applicability of optogenetics in mechanistic *in vitro* studies. We combine photo-uncaging or optogenetic neural stimulation with imaging of cardiac monolayers to measure electrical activity in an automated fashion, illustrating the power and high throughput capability of such interrogations. The methods described highlight the challenges and benefits of co-cultures as experimental model systems.

## INTRODUCTION

Cardiac impulse formation and conduction are modulated by autonomic activity, and the autonomic nervous system plays an important role in the initiation and maintenance of arrhythmias in diseased hearts [1, 2]. Sympathetic nerves release noradrenaline, which activates cardiac β-adrenergic receptors to modulate myocyte repolarisation and calcium handling via alterations of transmembrane currents and intracellular calcium homeostasis [3]. Increased sympathetic activity, which can occur during epileptic seizures [4] and is also associated with chronic diseases such as hypertension [5] and heart failure [6], is often associated with increased risk of re-entrant arrhythmias [7]. Tissue damage can also alter the distribution of innervation where cardiac cell death following myocardial infarction causes sympathetic denervation followed by nerve sprouting and reinnervation.

Nerve sprouting may promote the heterogeneity of excitability and refractoriness, which was suggested to increase arrhythmia susceptibility in the reinnervated infarct border zone [8, 9]. However, recent clinical studies [10, 11] have shown that cardiac sympathetic denervation (rather than reinnervation) can lead to a higher risk of ventricular arrhythmias and arrhythmic death. Experimental and computational studies linked the beneficial effect of innervation to attenuation of infarct-induced vulnerability to repolarisation alternans via β-adrenergic activation [12, 13] or to reduction of electrophysiological heterogeneity and calcium mishandling, which was present even when the nerves were not activated [14]. Resolving the unclear pro- or anti-arrhythmic effect of post-infarction reinnervation may also involve the precise understanding of the neural heterogeneity and its role in arrhythmia modulation. This type of research may benefit from the use of a cell culture model system where the effects of innervation can be precisely controlled. Co-cultures of cardiac myocytes and sympathetic neurons have been investigated for over 30 years [15, 16], however these studies were carried out at microscopic (single cell) scales where arrhythmogenicity cannot be directly assessed. Tissue heterogeneity and impulse conduction velocity (CV) play key roles in the initiation and stability of re-entrant spiral waves [1]. While CV depends in part on the excitability of individual myocytes, it also depends on cell-cell connectivity and tissue heterogeneity [17, 18].

Confluent myocyte monocultures imaged at macroscopic space scales have allowed the investigation of more complex functional tissue level properties such as wave propagation and pattern formation[19], and their ability to support reentrant spiral waves has validated their use as a model of arrhythmogenesis. Optical mapping of these cultures has given an insight into important arrhythmogenic mechanisms, including unidirectional conduction block, junctional coupling, and remodeling [19]. Confluent co-cultures of myocytes and neurons imaged at macroscopic scales (>1 cm^2^) are a potentially useful biological model system for the study of the proarrhythmic effects of abnormal sympathetic activation on cardiac conduction.

In this study, we report the first macroscopic optical mapping measurements of cardiac monolayers co-cultured with cardiac sympathetic stellate neurons imaged using our recently published dye-free optical imaging method [20]. We examine the physical contacts and test the functional coupling between cells in these co-cultures and make observations relating to changes in excitation patterns when neurons are pharmacologically stimulated. Finally, we explored the effects of neuron numbers on cardiac behavior and their ability to modulate cardiac excitability and found that only a limited number of neurons (one neuron connected to its neighbors in a low neuron to myocyte ratio) capable of driving a cardiac syncytium.

## RESULTS

### 1] Stellate Sympathetic Neurons make contacts with cardiomyocytes

Scanning electron microscopy of sympathetic neurons growing in co-culture with cardiomyocytes *in vitro* shows connections between neurite extension and cardiac syncytium (Figure 1 A). The neuron bodies and extensions clearly make physical contact with myocytes (Figure 1 B, C, D, E, F, G, H, I, J). Immuno staining of co-cultures shows co-localisation of Synpasin and Beta2 receptors (Figure 1 K), sympathetic neurons show positive staining for Th (Figure 1 L), and fibronectin contamination in the co-cultures was assessed by staining with Vimentin (Figure 1 M), which showed low abundance.

**Figure 1:**
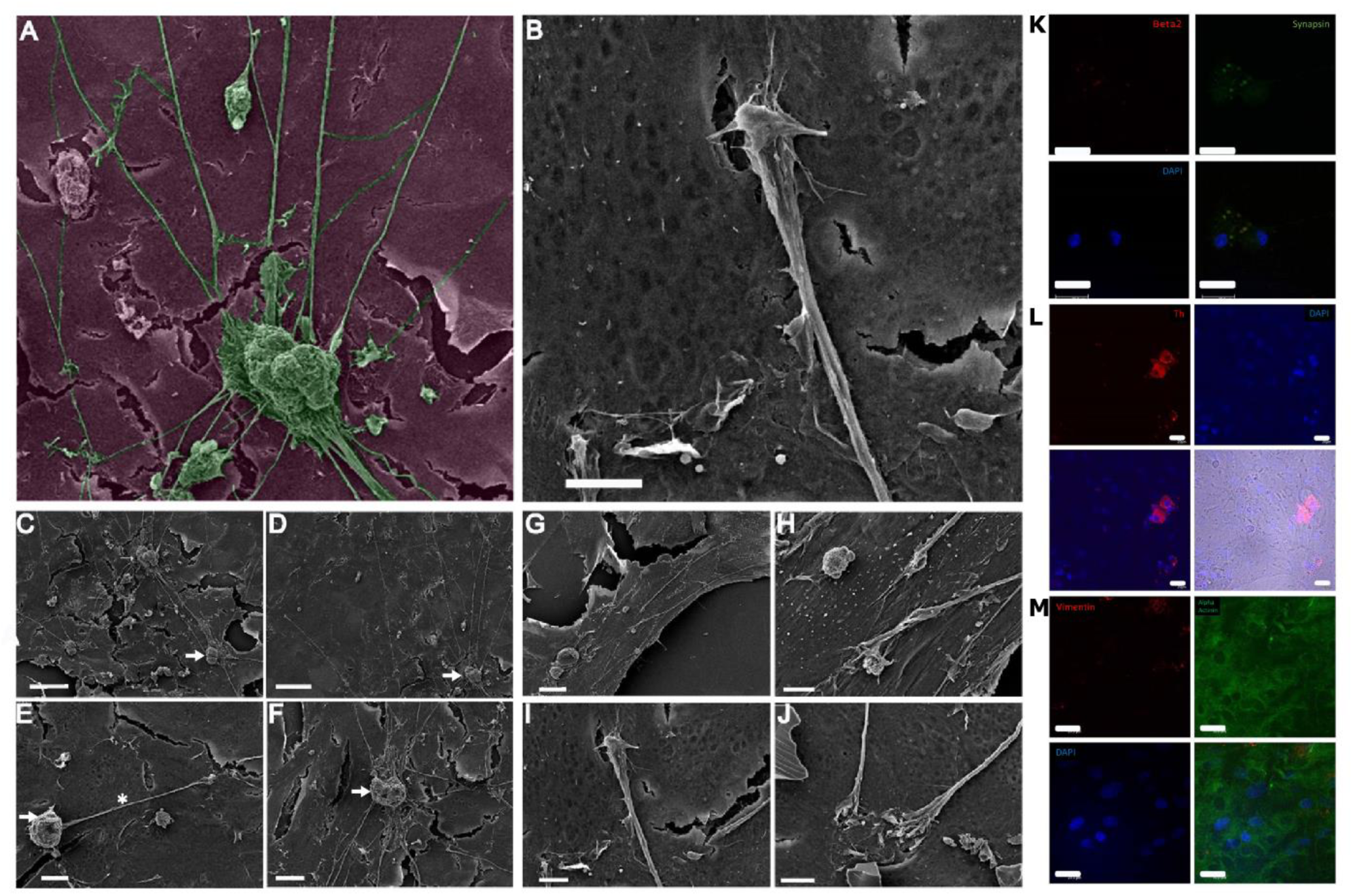
Scanning electron microscopy of cardiac-neuron co-cultures. (A) Exemplar image for sympathetic neurons growing in co-culture with cardiomyocytes in vitro (micrographs were colored in post-processing). (B) Close up of a connection between neurite extension and cardiac syncytium showing connections between neurons and myocytes. Scale bar 5 µm. (C-F) Neuron bodies and extensions make contact with myocytes, white asterisk showing axon and white arrow indicating neuron body. Scale bars 50, 50, 10 and 20 µm respectively. (G-J) Images of axon cones making connections with cardiomyocytes. Scale bars 20, 5, 5 and 5 µm respectively. (K) Immunohistochemistry of co-cultures labeled with Beta-2 adrenoreceptor and synapsin antibodies, (L) Th and DAPI and (M) Vimentin and alpha actinin co-staining on co-cultures. Scale bars 20 µm.

### 2] Pattern formation is affected by the presence of neurons

Using dye-free imaging (Figure 2 A, B, C), we investigated how neuronal activation modulates cardiac patterns of activation in monolayer culture. We chose experimental conditions that spontaneously yield a wide variety of wavefront topologies within the imaging system’s 16 × 16 mm field of view. Introduction of an additional cell type can potentially introduce heterogeneities that would impact wave front stability. Surprisingly, co-cultures displayed fewer wave breaks than their monoculture counterparts at similar plating densities (Figure 2 D). We broadly classified wave dynamics as simple (periodic target waves and single spiral wave reentry, Figure 2 D top row), or complex (single dominant spirals with additional irregular waves and multiple equally sized wavelets, Figure 2 D bottom row). While monocultures frequently displayed complex dynamics, co-cultures rarely displayed this behavior (Figure 2 E, P<0.05, chi-square test). Indeed, we observed wavelet reentry in only one of the co-culture preparations (6 isolations, 20 preparations).

**Figure 2:**
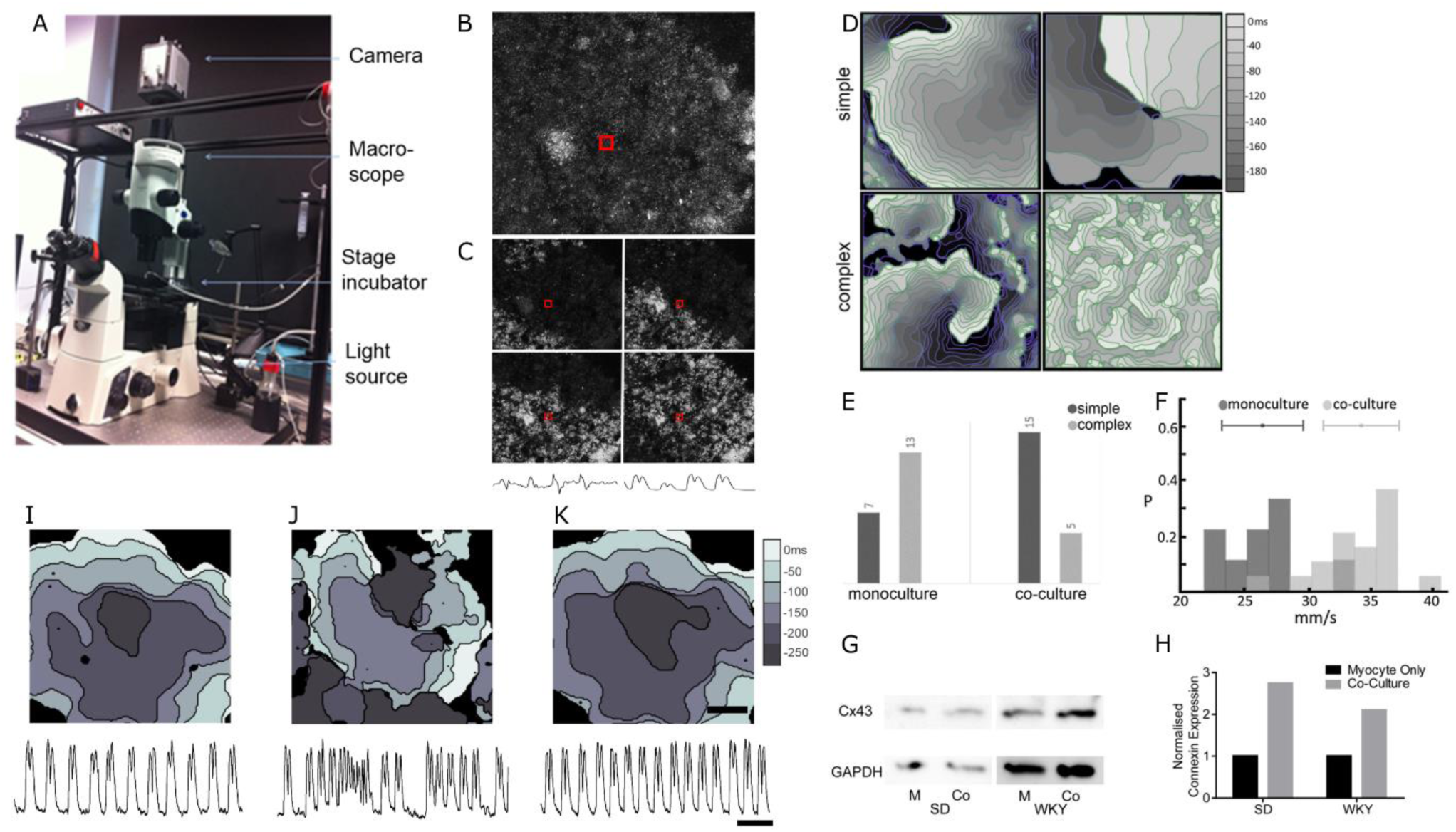
(A) The imaging setup uses off-axis illumination to obtain high contrast images of the monolayer. These images are processed to calculate the absolute value of the intensity change over n frames. (B) Unfiltered image (Note: The image is obtained using a 10x lens instead of the 1x used for the main dye free experiments). (C) Filter applied to track motion in the cardiac monolayer, below, intensity vs time plots for the central 5×5 pixels in the image [Filter: P_t_(x,y)-P_t_-N(x,y)]. The left trace shows intensity from unprocessed images and the right trace show the improved s/n after processing. (D) Wave dynamics in monocultures and co-cultures. Isochronal maps of wave dynamics in confluent cardiac-stellate neuron co-cultures display a variety of complex rhythms similar to those seen in intact hearts. Wave dynamics here are classified as simple (targets or single spirals) or complex (multiple spiral waves or wavelets of activity). (E) Monocultures display more complex dynamics than co-cultures, which display predominantly simple wavefronts with few wavebreaks (P<0.05, Chi-square). (F) Histogram representing the 90-percentile of wave speed from monocultures and co-cultures. (G and H) Western blots of Connexin43 expression in cultures grown in the absence and presence of sympathetic cardiac stellate neurons in two separate co-culture experiments (SD and WKY). (I-K) Nicotine stimulation causes a reversible change in the activation pattern. Top panels show isochronal maps, and bottom panels show binned intensity vs. time plots for the four center pixels. (I) A target wave before nicotine addition; (J) Irregular reentrant activation after addition of nicotine; (K) Washout allows the co-culture to revert to a target wave. See Supplementary Video 1 ppt.

### 3] Conduction velocity in spontaneously active co-cultures is faster than myocyte monocultures

We measured conduction velocity in unstimulated co-cultures (n=19) and myocyte monocultures (n=9). Figure 2 F shows a histogram representing the 90-percentile of wave speed from each recording. The conduction velocity in the co-cultures was significantly faster than in myocyte monocultures. Overall, the mean conduction velocity (± stdev) in the myocyte monocultures was 27 (± 3) mm/s and 34 (± 3) in the co-cultures. To try and understand the molecular mechanisms, we conducted a qualitative label free proteomics screen on the myocyte and co-cultures (Supplementary Tables 1 and 2). We found changes in pathways regulating gap junction protein expression along numerous changes in pathways associated with metabolism and development (Supplementary Figure S1). Additional measurements of Connexin43 (Cx43) levels in cultures using Western blot technique (Figure 2 G and H) confirmed that Cx43 was higher in the co-cultures (two independent experiments). Interestingly, in different cardiac co-cultures that are well connected and not spontaneously active/arrhythmic, the stimulated (1 Hz) conduction velocities for cultures with different neuron concentrations were very similar at 1Hz electrical pacing (Supplementary Figure S2), showing no significant difference using ANOVA followed by Tukey-Kramer.

### 4a] Nicotine stimulation can cause a transition between target and reentrant waves

In 8/12 cases, nicotinic stimulation, which is an established method of activating sympathetic neurons, did not markedly alter the pattern of activation in co-cultures. However, in four cases we observed changes in spatiotemporal patterns of activation following nicotine stimulation. Figure 2 I-K shows isochronal activation maps during application and washout of nicotine. The co-culture displayed target waves prior to nicotine application (Figure 2 I), which transitioned to reentrant spiral wave activity shortly after addition of nicotine (Fig 2 J). Washout of nicotine abolished the spiral wave (Figure 2 K), see Supplementary PPT Video 1 for corresponding raw data video file. Also see online video V1a,b and V2a,b showing pre-nicotine behavior (a) and post-nicotine behavior (b) in two separate experiments.

### 4b] Nicotine stimulation increases beat rate in co-cultures displaying target patterns

To confirm the formation of functional connections between cardiac myocytes and sympathetic stellate neurons, we stimulated the neurons in co-cultures with nicotine (n=6), following the protocol described in [21]. In co-cultures displaying target patterns we observed an average increase in beat rate of 31%±10 when 10 µM of nicotine was added (5 min post nicotine addition, and then the beat rate returned to baseline (Figure S3)). We also performed: (i) control experiments in which nicotine was added to cardiac monocultures (i.e. no neurons present) and we did not see any changes in the contraction rate; (ii) vehicle experiments (sterile distilled water) in co-cultures, which also showed no increase in beat rate.

### 4c] Bath application of nicotine produces a marked increase in spontaneous firing of action potentials in co-cultured neurons

Using the whole-cell current clamp method we observed repeated membrane depolarization events (in three separate experiments) on the addition of 10 µM nicotine (Figure 3 A). We measured the membrane potential of individual neurons within the co-culture at 0 – 60 s (pre) to 300 – 360 s (post) nicotine application, and found a significant rise in membrane potential (P<0.05, paired t-test). Figure 3 B shows representative video analysis of traces (performed simultaneously during patch clamp) from one of the experiments where macroscopic rate changes can be observed upon nicotine addition to the co-culture.

**Figure 3:**
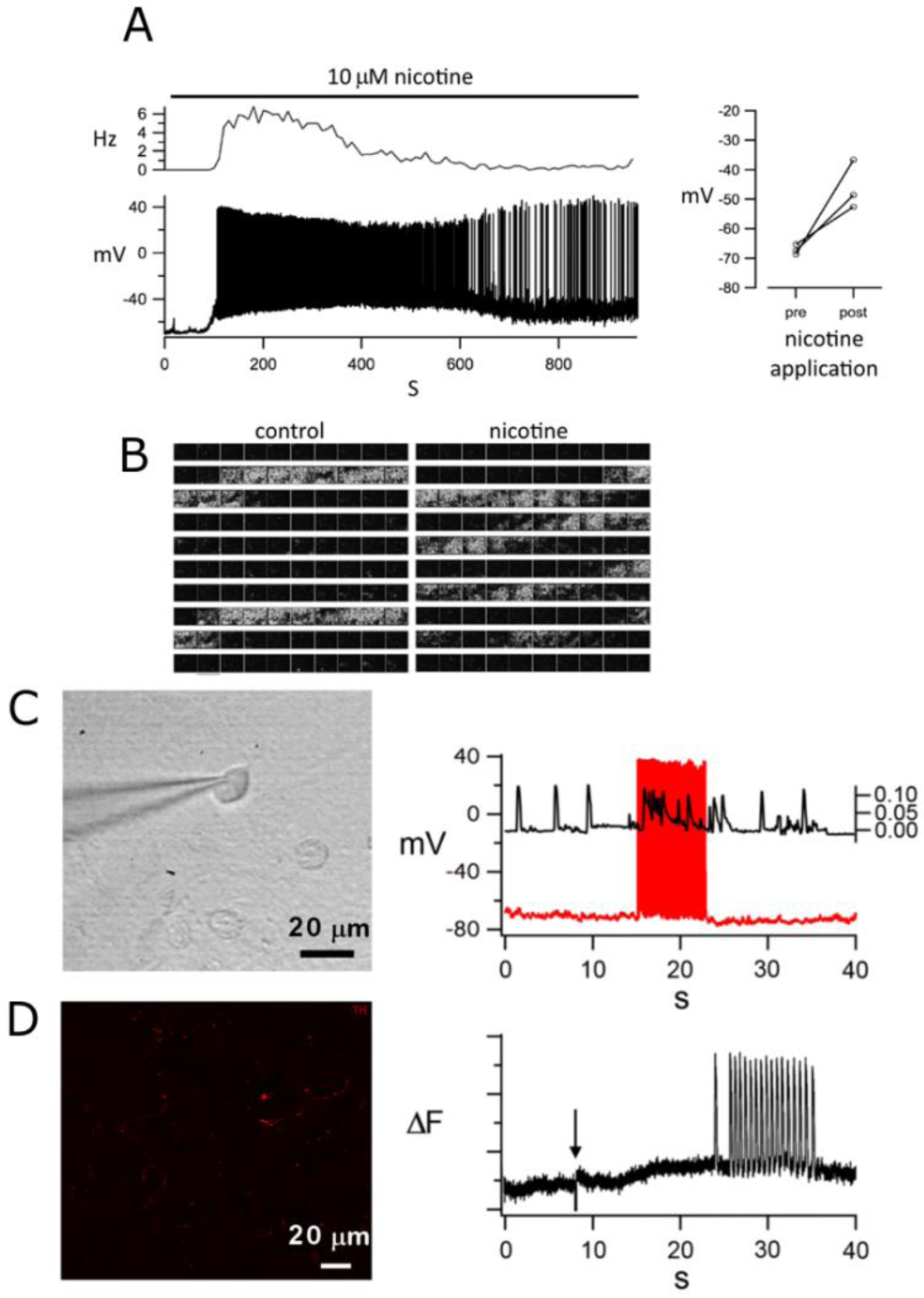
Simultaneous electrophysiological measurement of neuron membrane potential and video imaging of cardiac dynamics. (A) Addition of nicotine to co-cultures causes an increase in the rate of spontaneous firing in a patched neuron (p<0.05, paired t-test). The side panel shows the net depolarization of the resting potential of neurons before and after nicotine application. (B) Individual frames from a video recorded before and after nicotine application. The video was processed to show motion transients (white) as described in the supplement. (C) A single neuron modulates the activity of many connected cardiomyocytes. Current injection into a patched neuron causes rapid activation of the neuron (red trace), which is correlated to irregular activity in the monolayer (black trace), using frame cross-correlation in order to map changes. Synchronization of video imaging and electrophysiological trace is accurate to +/-2s. (D) OptoDyCE imaging of nicotine stimulation of a co-culture with a low density of neurons (1 neuron: 100,000 myocytes) results in increased activity in the monolayer (arrow-indicates uncaging of nicotine using blue light, n=12/18 responders). Red fluorescence indicates Tyrosine hydroxylase positive (Th) immuno staining specific to sympathetic neurons.

### 5] One neuron potentially stimulates a connected monolayer of one hundred thousand myocytes

Pilot electrophysiology experiments suggested that a single neuron could potentially modulate the activity of many connected cardiomyocytes. (Figure 3 C). Current injection into a patched neuron caused a rapid activation of the neuron, which was correlated to irregular activity in the monolayer by calculating the cross-correlation between sequential frames in order to map changes. Electrophysiological measurement on neuron activity in co-cultures is challenging as the underlying cardiac tissue moves with every beat resulting in lost patches. We therefore utilized an all-optical approach designed for high throughput interrogation of cardiac preparations namely OptoDyCE [22] to address this question and used optical uncaging of nicotine to study neuronal stimulation of myocyte behavior (Figure 3 D).

### 6] Optogenetic and Optochemical neuronal stimulation and the effect of neuro transmitter release on cardiomyocytes using OptoDyce

Co-cultures with different neuron concentration were plated in small (6.2mm diameter) wells at plating density of 140,000 myocytes / well. The stellate sympathetic neurons were infected with hChR2-eYFP. Schematic representation of the co-cultures with different neuron dosing regimes (neuron-myocyte ratios in 1:5, 1:20, 1:100, 1:100,000) are shown in Figure 4 A. Immunohistochemistry was used to confirm the presence of sympathetic neurons (using Th antibody) in the different neuron-myocyte co-culture ratio combinations (Figure 4 J). OptoDyce (all-optical dynamic cardiac electrophysiology framework) high throughput co-cultures also allowed the study of the effects of sympathetic cardiac stellate neurons on cardiac activity in well-connected quiescent cardiac cultures using optical mapping (cultures loaded with dye Di-4-ANBDQBS). Co-cultures were found to be constitutively more active than monocultures of myocytes (Figure S4). Fishers exact test (two sided), p= 0.0046 statistically significant. n=6 myocytes and n=24 for co-cultures.

**Figure 4:**
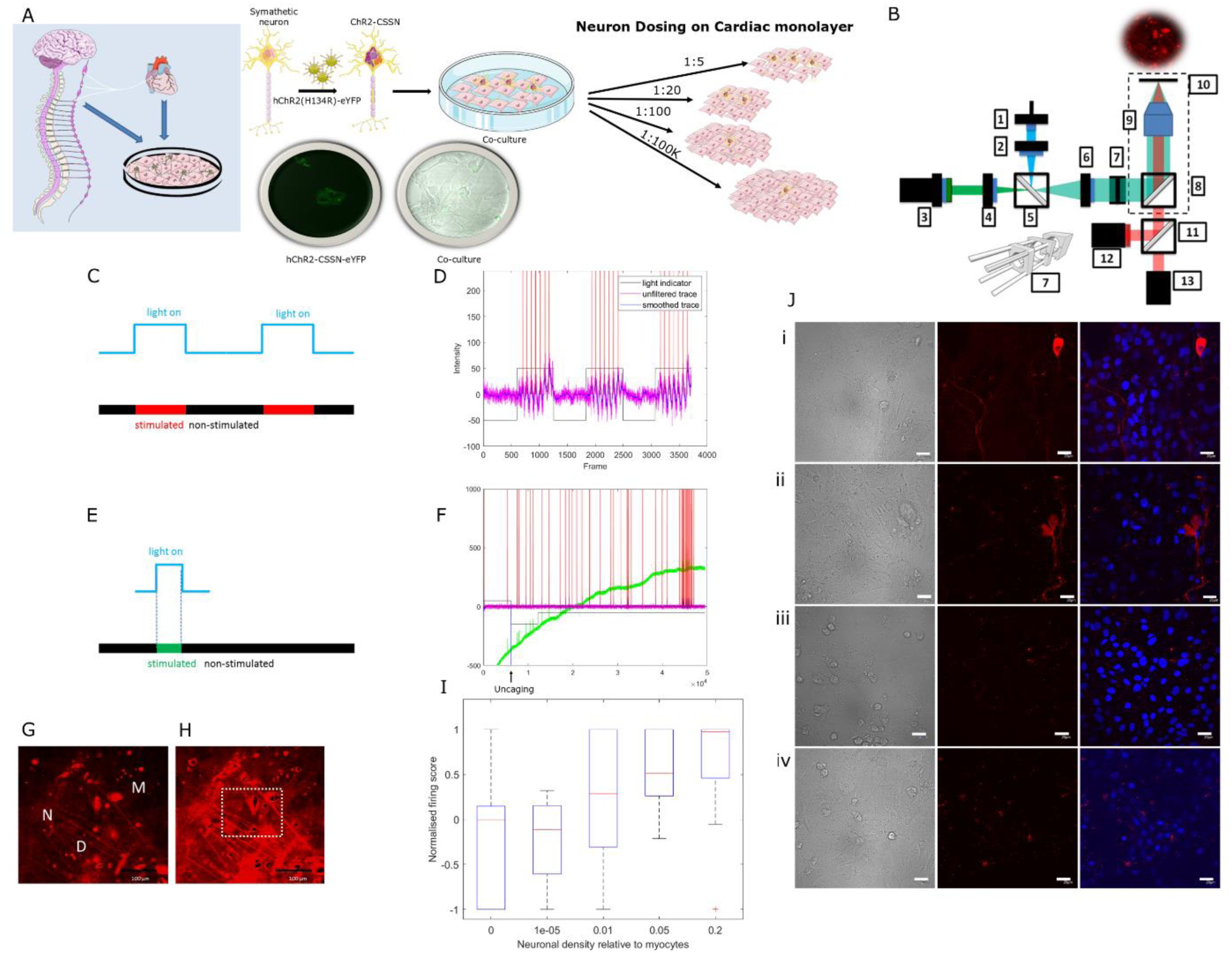
(A) Schematic representation of the *in vitro* co-culture system, neonatal rat cardiac stellate sympathetic neurons and neonatal rat ventricular cardiac confluent monolayers cultured to test sympathetic-cardiac interactions. The stellate sympathetic neurons were infected with hChR2-eYFP. Schematic representation of the co-cultures with different neuron dosing regimes (neuron-myocyte ratios in 1:5, 1:20, 1:100, 1:100,000). (B) OptoDyce set up: High throughput all-optical electrophysiology OptoDyCE with 96-well format, custom-built optical setup and automated protocol[22]. Fully automated OptoDyCE optical setup. Where 1 – LS1; 2 –L1; 3 – LS2 with L2 and F1; 4 – L3; 5 – DM1; 6 – L3; 7 – Adapter; 8 – DM2; 9 – Obj; 10 – Sample; 11 – DM3; 12 - D1; 13 - D2 with F2. (C and D) Illustration of the experimental protocols for “opto-electric” vs “opto-chemical” stimulation of neurons. (C) Optogenetic neural stimulation of cardiac tissue via Channelrhodopsin2 (ChR2), selectively expressed only in the neurons. Co-cultures of neurons and myocytes (loaded with dye Di-4-ANBDQBS spectrally compatible with ChR2) prior to stimulation (G) and after light activation (H). Optical stimulation (470 nm) was provided at pulse lengths of 5–20 ms, at 0.5–8 Hz, using irradiances of 0.4–7 mW mm−2, as needed. (D) Post processed traces using custom-written Matlab software. Traces showing baseline no activity and followed by long light pulse stimulation, action potentials are evoked indirectly in the myocytes via the ChR2-light-sensitized neurons. Magenta is the raw trace, blue is the trace after baseline subtraction after median filtering, red indicates detected spike times, black is an indicator of when light is present (black down=light off). N=neuron, D=dendrites, M=myocyte in (G). (I) The number of neurons innervating the myocytes affects the firing frequency of myocytes in cultures with different neuron to myocyte ratios (0.2=1:5, 0.05=1:20, 0.01=1:100, 1e-05=1:100,000 and 0= myocytes/controls); the number of experiments (n) for each group were 14, 4, 23, 28, 12 and confidence values (p) (against null hypothesis of zero effect; Wilcoxon signed rank test) were 0.7131, 0.6250, 0.1091, 0.0000 and 0.0352. We observe a dose-dependent effect (i.e. the greater the number of neurons innervating the myocytes, the greater the effect, with values greater than 0 indicating an increase in beat rate). (E-F) Uncaging of nicotine using a flash of blue light may lead to the release of noradrenaline by the sympathetic neurons resulting in increase in myocyte beat rate. (J) Immunohistochemistry analyses using TH (red) antibody in the different neuron-myocyte co-culture ratio combinations ((i) 1:5, (ii) 1:20, (iii) 1:100 and (iv) 1:100,000).

Figure 4 B shows a schematic of the high-throughput all-optical electrophysiology system OptoDyCE. Optogenetic neural stimulation of cardiac tissue via Channelrhodopsin2 (ChR2), selectively expressed only in the neurons was performed using a light stimulation protocol schematically represented in Figure 4 C. Co-cultures of neurons and myocytes were loaded with dye Di-4-ANBDQBS (Figure 4 G, H), which is spectrally compatible with ChR2. Fluorescence traces before and after light stimulation (Figure 4 D and Figure S5). Long light pulse stimulation results in action potentials evoked indirectly in the myocytes by ChR2-expressing neurons (Figure 4 D). Cardiac response to light stimulation of ChR2-expressing neurons show a dose dependent effect (Figure 4 I), where cultures with higher neuron concentrations generate more cardiac activity with the same light stimulus.

We also assessed using an optochemical approach where caged nicotine was added to each well (Figure 4 E and F), and nicotine is released with light stimulation (Figure S6). While we observed cases where low-density co-cultures responded to nicotine uncaging (12/18 responders), either by increasing beat rate or by inducing bursting behavior in the cardiac monolayer, these responses occurred after a long delay which raised the possibility that the effects are due to chance. Longer recording times both pre and post light stimulation are required to confirm the efficacy of this method.

Standard nicotine treatment of co-cultures with nicotine to drive sympathetic neurons offers very little spatio-temporal control over the experiments (Figure S7). Optochemical methods to cause uncaging of nicotine to stimulate neurons which in turn stimulate myocytes can be achieved (Figure 4 F and Figure S6). The timing of chemical release to stimulate neurons in culture and observe the myocyte response (rate) can be controlled with some precision (Figure 4 E). At the same time, optogenetic stimulation of stellate sympathetic neurons offers far superior precise spatio-temporal control of neuron behaviour and their effects on myocytes (Figure 4 D and Figure S5).

Additionally, to show the potential of applying all optical interrogation to hiPSc-derived cardiomyocyte-neuronal co-cultures, we performed several qualitative experiments involving optogenetics (driving hiPSc-derived peripheral neurons with a primary concentration of sympathetic neurons purchased from Axiogenesis (now Ncardia)). We show similar light control experiments to the neonatal co-culture experiments (See Figure S8).

## DISCUSSION

Direct neural-cardiac interactions (at the cell level) are an understudied area due to lack of specific tools with high spatio-temporal resolution. We demonstrate that interesting questions like the effect of neural density on the electrophysiological properties of a cardiac syncytium can be studied in a high-throughput manner using all-optical techniques. Photo-uncaging of nicotine or optogenetic neural stimulation are used here in conjunction with optical imaging of cardiomyocyte contractile and electrical activity to illustrate the power of such interrogations. We designed a simplified *in-vitro* model of neurally modulated arrhythmogenesis by co-culturing stellate sympathetic neurons with confluent monolayers of myocytes, and optically measured the effects of these neurons on cardiac wave speeds using a newly developed dye free imaging method. We demonstrate that physical (Figure 1) and functional connections (Figure 2–4) are formed between cardiac sympathetic stellate nerve cells and cardiomyocytes as previously reported in isolated cell preparations [23, 24]. Novel findings herein are that (i) neurons have a significant impact on base wave speed when cultured with spontaneously arrhythmic cardiomyocytes and no significant effects on conduction velocity in well-connected cardiac monolayers; (ii) In our culture model, co-cultures display a higher degree of spatial organization: while myocyte monocultures frequently display multiple interacting wavelets of activity, co-cultures are more likely to be driven by a single dominant spiral (Figure 2). In addition, we are able to observe transitions between target waves and re-entrant dynamics as a consequence of neural activation, and demonstrate that increased spike frequency in neurons correlates with increased cardiac activity at macroscopic scales. Finally, we used a high throughput voltage sensitive dye based imaging modality to show that the number of neurons needed to modulate cardiac dynamics is important.

While our experiments demonstrate that innervation results in functional changes in CV and the spatial organization of waves in culture, we do not know the mechanisms responsible for these changes. Fibroblast concentration, gap junction density and ion channel expression are known modulators of conduction velocity. Immunofluorescence studies showed no evidence of increased vimentin expression indicating low fibroblast proliferation (Figure 1 M), however, gap junction protein Cx43 was elevated in co-cultures relative to cardiac monocultures (Figure 2 G, H). Additionally, we performed a label-free quantitative proteomics screen on cultures (See Supplement Table 1 and 2). We found changes in pathways regulating gap junction protein expression along numerous changes in pathways associated with metabolism and development (Supplement Figure S1). The results of our proteomics screen is consistent with other studies which link innervation to developmental processes [21, 25–28]. Shortly after birth, cardiomyocyte hyperplasia decreases and CV increases [25], which correlate to increased β-adrenoreceptor density on cardiomyocytes and higher levels of catecholamines in the circulation [29]. In 2015 [30], experiments demonstrated that having sympathetic neurons present in *in-vitro* cardiac cultures delays cardiomyocyte cell cycle withdrawal and transiently limits hypertrophy via a β-adrenergic signaling pathway, which suggests that sympathetic innervation can regulate cardiomyocyte numbers during the postnatal period. Developmental changes may occur in neurons as well: Oh et al 2016 reported increased maturation of hiPSC-derived sympathetic neurons in their cardiac neuron co-culture system [31]. Coppen et al. have suggested the existence of post-natal changes in connexin expression in the developing fetal heart [32]. There have been other suggestions that expression of different connexin isotypes varies not only within distinct compartments of the adult heart, but also as a function of cardiac developmental stage[33]. It is possible that neurons enhance maturity of cardiomyocytes and the CV increase seen in our experiments may be due to developmental changes in the cardiac myocytes and improved connectivity.

Furthermore, our tissue culture results may also be relevant to understanding the effects of nerve sprouting in scar tissue, and in helping to resolve the apparently conflicting results summarized in the Introduction. Our observation that co-cultures display fewer wavebreaks than monocultures, have increased CV and higher levels of Cx43 offer indirect support for the protective role of neurons in intact tissues, particularly in cases such as the infarct border zone, which shows reduced function of gap junctions [34]. At the same time, the effects of acute nicotinic stimulation of neurons in our co-culture system may be a model of proarrhythmogenicity in the hyperinnervated infarct border zone.

Whilst conventional electrophysiology techniques allow for specific and micro control of single or sparse cell cultures, the application of such techniques is technically challenging when it comes to studying two cell types grown in a syncytium (such as the spontaneously excitable cardiac tissue and neurons). As was hypothesised may years ago by Arthur Winfree [1], the pattern of nervous system innervation could determine whether an arrhythmia could be instigated. Alterations in autonomic function occur in several interrelated cardiac conditions including SCD, congestive heart failure, diabetic neuropathy, and myocardial ischemia [35]. Neural modulation as a treatment for arrhythmias has been well established in certain diseases (such as long QT syndrome), however, in most other arrhythmias, it is still an open question and the subject of intense research [36]. Ongoing research over the last five decades has highlighted the importance of communication between neural and cardiac tissues. While experimental challenges still need to be overcome, dissecting mechanisms along the heart-brain axis has become more achievable with the introduction of novel imaging [17,19] and tissue engineering techniques [34].

The utility and scope of our macroscopic co-culture model offers even greater potential. In addition to using dye free approaches [20] to measure pattern formation and conduction velocity, the OptoDyCE technique [22] can be used for high resolution, high-throughput interrogation of neural influence on cardiac monolayers using optochemical and optogenetic stimulation (Figure 4). We have also extended this line of investigation from neonatal cells to Human iPSc-derived cardiomyocytes and iPSc-derived peripheral neuron co-cultures as a proof of concept study (Axiogenesis (now Ncardia), Supplement Figure S8). Thus, the methods described here provide approaches that could broaden our insight into fundamental disease mechanisms. The use of *in vitro* techniques in pharmacological assays and profiling is growing in its popularity in the drug discovery process [37]. Our experimental model in conjunction with recently developed imaging platforms can be applied to improving the efficacy of preclinical drug toxicity and discovery studies.

## MATERIALS AND METHODS

### 1] Spontaneously active cell culture

Here we use a neonatal ventricular cardiac monolayer cell culture model that spontaneously displays a wide range of behaviors [20] to investigate how neurons modulate pacemaking and reentrant activity. We measured activity (i) using a macroscopic dye free optical mapping imaging modality, (ii) patch clamp electrophysiology coupled with video microscopy and (iii) microscopic optical mapping. All experiments were performed in accordance to UK Home Office Animals Scientific Procedures Act (1986). Hearts were isolated from neonatal SD rat pups (P1-P3), killed by Schedule 1 in accordance to UK Home Office Animals Scientific Procedures Act (1986). Ventricular myocytes were enzymatically isolated by a series of enzymatic digestions in trypsin (1mg/mL, Sigma Aldrich, UK) followed by collagenase (1mg/mL, Sigma Aldrich, UK) and triturated to achieve a suspension of cardiomyocytes. The isolated cells were then pre-plated in an incubator (37°C, 5% CO2) for an hour to allow fibroblasts to settle at the bottom of the dish. The ventricular myocytes in the supernatant were then carefully removed from the dish and a cell count performed using a haemocytometer and trypan blue. The myocytes were plated on 35 mm poly-lysine coated petri-dishes (Bio coat Poly-D-Lysine 35mm petri-plates, Corning, UK) at a density of 750,000 cells (per 35 mm petri-dish) in plating medium (85% DMEM, 17% M199, 10% Horse serum, 5% FBS and 1% penicillin/streptomycin, all from Sigma Aldrich). 24 hours later the cardiac sympathetic stellate neurons were isolated from litter mates as described previously with some modifications [38, 39]. Briefly, following microscopic dissection of the sympathetic stellate ganglia, enzymatic digestion in trypsin 1mg/mL (Worthington, USA) and collagenase type-4 1 mg/mL (Worthington, USA), cells were dissociated by sequential mechanical trituration using fine (fire-polished) glass pipettes. Neurons were pre-plated for 1 hour (to eliminate fibroblast and Schwann cells), counted and plated in at varying neuron to myocyte ratios. Co-cultures were created by plating neurons on top of the cardiac monolayers. Co-cultures were maintained in media supplemented with nerve growth factor (50 ng/mL, NGF Millipore), which promotes neuron development. The same high serum level media was refreshed every other day, which supports the spontaneously active cardiac monolayers.

#### Drugs

Stimulation of cardiac sympathetic stellate neurons with nicotine: To measure the effects of neuron stimulation on myocyte beat rate, 10 µM nicotine ([−] nicotine hydrogen tartrate salt; Sigma-Aldrich) was used. Washout was performed with pre-warmed media. In case of caged nicotine experiments, caged nicotine known as RuBiNic (RuBi-Nicotine, Cat No. 3855, Tocris) has been used in neurotransmission studies [40]. Uncaging can be extremely rapid, controlled in time or space and quantitatively controlled and repeated. We tested the following neuron-myocyte densities: 1:5, 1:20 and 1:100. We employed caged nicotine at 100 µM as the actual amount of uncaged drug is usually extremely small based on details in [41].

### 2] Dye-free measurement of wave dynamics

We used dye-free imaging techniques [42] with modifications as described in [20] to image the spontaneously active confluent monolayers. Here, we employed an Olympus MVX10 Macroscope and Andor Neo sCMOS camera to record wave patterns and beat rate (from day 3 onwards). Experiments were carried out in an Okolab (Indigo Scientific, UK) stage incubation chamber controlled for heat (33-37°C), CO2 (5%) and humidity. Culture plates were allowed to equilibrate in these conditions for 20 minutes before commencing recordings. From these we were able to observe the propagation of cardiac waves across the plates by detecting the minute contractile motions of the myocytes. Frame rates were captured between 50 to 100 fps depending on the desired record duration. The software for displaying the camera output in real time, saving and analyzing data were written in a combination of Java and Python, and is directly available from the authors. For detailed description of the methods see [20]. To measure the effects of neuron stimulation on myocyte beat rate, 10 µM nicotine ([−] nicotine hydrogen tartrate salt; Sigma-Aldrich) was used to trigger depolarisations. Washout was performed with pre-warmed media (similar to the methods described by [17]).

### 3] Simultaneous patch clamp electrophysiology-video recording

Current clamp was used to record electrophysiological traces from neurons, while video recording of cardiac behaviour was performed simultaneously using a CCD camera. The signal was recorded and analysed using custom-made procedures in Igor Pro (Wavemetrics). Image series were after processing the images so that the value of each pixel p at frame t, denoted p(t), is replaced by the absolute value of p(t) − p(t−5).

Current clamp recordings were performed with a Multiclamp 700B amplifier (Molecular Devices). Borosilicate glass pipettes were filled with an internal solution containing (in mM): 110 potassium-gluconate, 40 HEPES, 2 ATP-Mg, 0.3 GTP and 4 NaCl (pH 7.2–7.3; osmolarity 270–285 mOsmol). Standard artificial cerebrospinal fluid (ACSF) bubbled with 5% CO2 was used as the bath solution, containing (in mM): 126 NaCl, 3 KCl, 1.25 NaH2PO4, 2 MgSO4, 2 CaCl2, 26 NaHCO3, and 10 glucose, pH 7.2–7.4. All recordings were conducted at 37°C. Square pulses of current or current steps were administered to depolarise the neurons and elicit action potential firing. Data were low-pass filtered at 2 kHz and acquired at 5 kHz. Further information can be found in the Supplement.

### 4] Assessment of activity from co-cultures with low neuron density

We performed a pilot study where a single patched neuron, driven to rapidly fire by injection of current, modulated the activity of a confluent monolayer. While this pilot experiment suggested that a single rapidly firing neuron can drive the behavior of a macroscopic cardiac monolayer, this protocol cannot rule out network effects from unpatched neurons. However, performing patch experiments on co-cultures with very few neurons (e.g. theoretically calculated one neuron per monolayer) was not practical as it proved to be technically difficult to locate a neuron amongst thousands of cardiomyocytes within an acceptable time frame. We therefore opted to investigate the activity co-cultures with low neuron numbers (calculated for 1 neuron:100,000 myocytes) using opto-chemical stimulation and OptoDyCE imaging [22]. Since the low number of neurons increased variability, experiments were performed using multi-well plates to obtain sufficient statistical power. The use of multi-well plates necessitated the use of a modified experimental setup that differed from the one used to image activity in conventional petri plates as follows. Neonatal rat ventricular cardiomyocyte Culture: Neonatal (2–3-day old) Sprague–Dawley rats were killed by Schedule 1 and ventricular tissue was removed as per an approved Stony Brook University IACUC protocol. The ventricular tissue was digested in trypsin made up in in Hanks’ Balanced Salt Solution (1 mg/mL, US Biochemical, Cleveland, OH) overnight at 4°C. the tissue was serially digested using 1 mg/mL collagenase (Worthington Biomedical, Lakewood, NJ) in HBSS at 37 °C and pipetted into conical tubes and placed on ice. After centrifugation, cells were re-suspended in culture medium M199 (GIBCO) supplemented with 12 μM L-glutamine (GIBCO), 0.05 μg/mL penicillin-streptomycin (Mediatech Cellgro, Kansas City, MO), 0.2 μg/mL vitamin B12 (Sigma-Aldrich, St. Louis, MO), 10 mM HEPES (GIBCO), 3.5 mg/mL D-(+)-glucose (Sigma-Aldrich) and 10% fetal bovine serum, FBS (GIBCO). Fibroblasts were removed via a two-step pre-plating process, where the cell suspension was plated in a flask and incubated (37 °C, 5% CO2) for 45–60 min and switched to a new flask and the incubation repeated. Cardiomyocytes were then counted using a hemocytometer before plating in glass-bottom 96-well plates (*In Vitro* Scientific). 24 hours later the cardiac sympathetic stellate neurons were isolated from litter mates and co-cultured with the cardiomyocytes as described above in Cell Culture Methods. Low serum (2%) maintenance media was used from day 2. Optical recording of membrane voltage, Vm, was performed using the synthetic voltage-sensitive dye Di-4-ANBDQBS. 17.5 mM of stock solution in pure ethanol is diluted to 35 μM Tyrode’s solution. Cells are stained for 6 min in dye solution followed by a 6 min wash in fresh Tyrode’s. Finally, the wash solution is replaced with fresh Tyrode’s. Imaging was performed at >200 frames per second (fps) with 4 × 4 binning using NIS-Elements AR (Nikon Instruments; Melville, NY). RuBi-Nicotine uncaging was elicited by a 150 ms, 1200 mA flash (470nm). Neurons were stimulated using 100 µM caged nicotine (Tocris, USA)[43] instead of free nicotine in solution in order to minimize the time between sequential measurements in multiple wells. The concentration of free nicotine following photorelease in these experiments was less than the concentration of the caged nicotine used (100 µM) as photorelease efficiency is poor at physiologically tolerated light levels. We tested the effects of caged nicotine between 10-100 µM and found that 100 µM resulted in neuronal driven cardiac responses similar to those evoked by the pure nicotine compound. We note that low photorelease efficiencies have been reported with similar compounds: Macgregor et al 2007 report a 2% efficiency of photorelease with caged NAADP [41].

The geometry of the multi-well plate (deep well relative to the bottom surface area) is not compatible with the oblique illumination method used for dye-free imaging used in larger plates due to unwanted light reflections of the sides of the well. In addition, the advantages associated with dye-free imaging (long term recording with low phototoxicity) are less relevant for these experiments as activity in different wells are measured in rapid succession for relatively short time periods. We therefore opted to measure cardiac activity using voltage-sensitive dye Di-4-ANBDQBS (10 µM, supplied by Dr Leslie Loew, University of Connecticut). A standard epi-fluorescence configuration was used (530 nm excitation light, with a 20x objective (20 x Nikon CFO Super Plan Fluor).

The short time available for each recording and the microscopic field of view precludes measurement of CV and relatively subtle changes in patterns of activation. We therefore use experimental conditions that allow for an unambiguous assessment of the effects of nicotine on cardiac dynamics. For cell plating, 96-well glass bottom dishes (In Vitro Scientific) were coated with 50 μg/mL fibronectin (diluted in PBS), and incubated at 37 °C for at least 2 h before cell plating. Cells are plated in 10% FBS M199 media; on day 2, the media was replaced with 2% FBS M199 until the day of experiments and imaged at room temperature, which favors quiescent or slowly beating cultures [22]. The co-culture in each well can then be labeled as responsive or un-responsive to nicotine based on the induction of activity in normally quiescent cultures or a marked increase in beat rate in slowly beating cultures. Wells that display rapid or irregular activity before the addition of nicotine are excluded from subsequent analysis.

Each well is plated with 140,000 cardiomyocytes and seeded with neurons at a concentration of 1 neuron per 100,000 myocytes (theoretical aim). At this plating density, Poisson statistics predict that 24.6% of plates will have zero neurons, 34.5% will have one neuron, and 40.9% will have more than one neuron. The number of wells that respond to nicotine is used to determine the number of neurons needed to induce activity in a connected monolayer.

### 5] Measurement of conduction velocity in well-connected cultures with optical mapping

Cardiac myocytes were cultured using the methods described in [44] and grown in glass bottom 35mm poly-lysine-coated Petri dishes (bio coat poly-D-lysine, 35 mm Petri plates, Corning; n=6). Neurons were isolated using the same methods described above. Co-cultures were created using different myocyte-neuron concentrations (20:1 n=6, 100:1 n=4, 600K:1 n=6). Using conventional optical mapping methods, we measured conduction velocity in well-connected cardiac cultures and co-cultures utilizing the dye Rhod-4-AM (10 µM: AAT Bioquest, Sunnyvale, CA). Rhod-4 AM diluted in Tyrode’s solution containing the following in mM: NaCl, 135; MgCl_2_ 1; KCl, 5.4; CaCl_2_, 1.5; NaH_2_PO_4_, 0.33; glucose, 5; and HEPES 5 adjusted to pH 7.4 with NaOH. All experiments were conducted at room temperature. Details for the macroscopic imaging system used in this experiment has been described previously [45]. Data was spatially and temporally filtered, using the Bartlett and Savistsky-Golay filters, before being analyzed in custom-developed Matlab (Mathworks, Natick, MA) software.

### 6] Structural studies (confocal microscopy and scanning electron microscopy)

#### Scanning electron microscopy (SEM)

Co-cultures were prepared for SEM using a protocol adapted from [46]. Briefly, cells were grown on 13 mm glass coverslips and fixed in 2.5% glutaraldehyde, post-fixed in 1% osmium tetroxide at 4°C for 1 hr, taken through an ethanol dehydration series and then dried for 3 minutes with HMDS. Cover-slips were mounted onto SEM stubs with a conductive carbon backing, coated with approximately 8 nm gold and imaged on a JEOL-6390 SEM (Figure 1).

#### Confocal microscopy

Briefly, cells were grown on 12 mm Poly-D-Lysine/Laminin coated coverslips (BD biosciences, UK) and fixed in 4% paraformaldehyde (Pierce Cat#28906) Following fixation, the cultures were washed in PBS and blocked with 10% goat serum and 0.3% BSA, and permeabilised with 0.1% Triton X-100 for 1 hour at room temperature. Cells were labelled with anti-sacromeric alpha actinin (1:650 A7811, Abcam ab182136), anti-tyrosine hydroxylase (1:250 T1299, Sigma Aldrich), anti-beta2 adrenergic receptor (1:100, ab182136, Abcam, UK) and anti-Vimentin (1:250 ab92547, Abcam, UK) primary antibodies overnight at 4°C to confirm the presence of sympathetic neurons in co-cultures and to assess the growth of fibroblast in culture with Vimentin. Secondary fluorescent antibodies (Alexa 488 and 647, 1:1000) were then applied for 2 h at room temperature (controls were labelled with secondary antibody only to ensure specificity of labelling, Figure 1 K-M). Slides were mounted in Vectashield to reduce photobleaching of fluorescence. Cultures were imaged using confocal microscopy (NT confocal laser-scanning microscope, Leica Microsystems Germany).

### 7] Western blot studies

The lysate was heated to 95°C for 5 min (to denature the protein) and vortexed before 20 µg was loaded into a 4-12% Tris-Glycine gel. The gel was run for 150 minutes at 125 V. After electrophoretic transfer (2 hours, 40 V) to a membrane (nitrocellulose), the membrane was washed with Tris Buffered Saline (TBS: 50 mM Tris-CI, 150 mM NaCl, pH 7.6) and blocked for an hour using a 5% milk solution (in TBST: TBS with 0.1 % Tween20). The primary antibody (Connexin 43, AB1728, Millipore, UK; 1:200) in 5% milk solution was incubated for 60 min at room temperature on a rocker. The membrane was washed three times in TBST at intervals of 10 minutes and incubated with a horse radish peroxidase (HRP)-conjugated secondary antibody (anti-mouse IgG 1:10000, anti-rabbit IgG 1:10000; both Novus Biologicals) in TBST solution for 60 minutes, then subjected to three washes at 10 minute intervals using TBST. Next, the membrane was washed twice for 10 minutes in distilled water for to ensure complete removal of tween which could interfere with HRP reaction. The ECL+ kit (Perkin Elmer Western Lightning ECL Pro, NEL120001EA) was used and a series of exposures using photographic film was collected. This was repeated with a GAPDH antibody, the loading control, (1:2500, Abcam, ab181602). Optical densitometry of the western blots was conducted using ImageJ software.

#### Statistics

Details provided in each result and online methods. Data are presented as means +/-stdev. In all cases P<0.05 was considered to indicate a statistically significant difference. The chi-square statistic was used to assess differences in pattern formation. The Kogmogorov-Smirnov test was used for the automated measurement of wavefront speed, and the t-test was used to assess patch clamp results. ANOVA followed by Tukey-Kramer was used for well-connected mono and co-culture conduction velocity statistical tests.

## Supporting information

Supplemental Video-V1a_co_prenic

Supplemental Video-V1b_co_nic

Supplemental Video-V2a_prenic

Supplemental Video-V2b_co_nic

## Acknowledgements

GB acknowledges salary support from Medical Research Council. RABB is funded by a Sir Henry Dale Wellcome Trust and Royal Society Fellowship (109371/Z/15/Z) and acknowledges support from The Returning Carers’ Fund, Medical Sciences Division, University of Oxford. RABB is a Winston Churchill Fellow and received some travel support from the Winston Churchill Trust for part of this study. RABB is a Senior Research Fellow of at Linacre College. RAC is a post-doctoral scientist funded by the Wellcome Trust and Royal Society. JT acknowledges support from the EPSRC and Bakala Foundation. NH is a British Heart Foundation (BHF) Intermediate Fellow (FS/15/8/3115).

This study was funded by the BHF Centre of Research Excellence Oxford (GB) and the EPSRC (Developing Leaders Grant held by RABB) and the Wellcome Trust and Royal Society (RABB). NH and DJP acknowledge support from the BHF and this study was also supported by a BHF project grant (PG/11/6/28660) to DJP and NH.

We would like to thank Dr Claudia Juarez Molina, Dr Suhail Aslam and Bevin Gangadharan for technical help. We also thank Dr Samuel Bose for commenting on the manuscript and Dr’s Winbo and Montgomery for scientific discussions that were supported by the Colin Pillinger International Exchanges Award.

## SUPPLEMENTARY MATERIALS

### Cell culture

Hearts were isolated from neonatal SD rat pups (P1-P3), killed by Schedule 1 in accordance to UK Home Office Animals Scientific Procedures Act (1986). Ventricular myocytes were isolated and subjected to a series of enzymatic digestions in trypsin (1mg/mL, Sigma Aldrich, UK) followed by collagenase (1mg/mL, Sigma Aldrich, UK) and triturated to achieve a suspension of cardiomyocytes. The isolated cells were then pre-plated in an incubator (37°C, 5% CO_2_) for an hour to allow fibroblasts to settle at the bottom of the dish. The ventricular myocytes in the supernatant were then carefully removed from the dish and a cell count performed using a hemocytometer and trypan blue. The myocytes were plated on 35 mm poly-lysine coated petri-dishes (Bio coat Poly-D-Lysine 35mm petri-plates, Corning, UK) at a density of ~700,000 cells (per 35 mm petri-dish) in plating medium (85% DMEM, 17% M199, 10% Horse serum, 5% FBS and 1% penicillin/streptomycin, all from Sigma Aldrich). 24 hours later the cardiac stellate neurons were isolated from litter mates as described previously [28, 39] with some modifications. Briefly, following microscopic dissection of the sympathetic stellate ganglia, enzymatic digestion in trypsin 1mg/mL (Worthington, USA) and collagenase type-4 1 mg/mL (Worthington, USA), cells were dissociated by sequential mechanical trituration using fine (fire-polished) glass pipettes. Neurons were pre-plated for 1 hour (to eliminate fibroblast and Schwann cells), counted and plated in a ratio of 1 neuron : 20 myocytes. Co-cultures were maintained in media supplemented with nerve growth factor (50 ng/mL NGF), which promotes neuron development. Media was refreshed every other day.

### Stimulation of cardiac sympathetic stellate neurons with nicotine

To measure the effects of neuron stimulation on myocyte beat rate, 10 µM nicotine ([−] nicotine hydrogen tartrate salt; Sigma-Aldrich) was used. Washout was performed with pre-warmed media.

### 98 well microscopic optical recording of co-cultures with voltage-sensitive dye using OptoDyCE method

Isolated neonatal cardiac cells were plated on fibronectin coated 96-well glass-bottom plates (In Vitro Scientific, P96-1-N). After 24 hours, stellate ganglia were isolated from litter mate pups (P3) and neurons isolated as described above, and plated on top of the myocytes. We performed optical recording of membrane voltage using the voltage-sensitive dye Di-4-ANBDQBS (from Leslie Loew, University of Connecticut), in normal Tyrode solution (4-5 days after cell plating). We compared a high dose versus low dose myocyte neuron co-culture combination (i.e. 1:5 (1 neuron stimulating 5 myocytes) and 1:100,000 (1 neuron stimulating 100,000 myocytes)) to determine the effect of neuron density on myocyte activity. Neurons were stimulated using 100 µM caged nicotine (RuBi-Nicotine, Cat No. 3855, Tocris). Measurements were carried out at room temperature as the multi-well plate setup did not accommodate a stage top incubator. We only analysed data from wells where the tissue was quiescent or beating with a rate lower than 1 Hz before nicotine stimulation: in particular recordings from monolayers that displayed bursting dynamics before nicotine stimulation were not used. Three of the wells displayed bursting dynamics before nicotine stimulation and were excluded from subsequent analysis.

### Structural Studies

Scanning Electron microscopy was carried out to observe the patterns of connections displayed between the sympathetic neurons growing in co-culture with cardiomyocytes *in vitro*. Immunofluorescence was performed on the cardiac-neuron cultures as follows: Beta-2 adrenoreceptor and synapsin antibodies (Figure 1 K), (Figure 1 L) Th and DAPI and (Figure 1 M) Vimentin and alpha actinin co-staining on co-cultures.

### Measurement conditions

Our experimental goal was to determine how the addition of neurons modulates activation patterns in cardiac culture under a wide range of initial conditions. We found that cultures plated on poly-D-lysine coated plastic petri dishes and continuously maintained high serum conditions (10% Horse serum, 5% FBS) resulted in isotropic cultures that spontaneously display a wide range of excitation patterns. In addition, we performed imaging experiments in maintenance medium gassed at 5% CO_2_ in a stage-top incubator (Okolabs) between 32°C and 37°C. Experimental conditions between research groups differ, but typically cultures by other groups are prepared by plating tissue on fibronectin coated glass coverslips, use low serum conditions after two days in culture, and perform imaging experiments after transferring dishes to a chamber in standard Tyrode solution in normal atmosphere. However, in our hands cultures prepared in these conditions often require pacing to induce activity, which was not compatible with our experimental aim of finding how the addition of neurons modulate spontaneous activity.

### Imaging setup

We developed a dye free imaging system similar to that described in [42], where off-axis oblique illumination is used to generate high contrast images that can be analysed to extract wave activity. We employ an Olympus MVX10 macroscope to relay a 1x image to a Neo sCMOS camera running at 50 frames/second. The high resolution of the camera allows visualization of the contraction of the tissue at the cell level, while still giving a relatively large 16 × 16 mm field of view. High resolution frames are processed so each displayed frame is generated by subtracting a frame captured at t-300 ms from the current frame and displaying the absolute value of each pixel. Intensity vs. time traces are obtained by 5×5 pixels in the image. The intensity vs. time traces typically have a double hump morphology which is due to contraction (first hump) followed by relaxation (second hump) of the tissue.

### Wave speed measurement

We employed an automated approach for finding cardiac waves in optical mapping recordings and measuring their speed of propagation. To compare wave speed between different co-cultures and myocyte monocultures, we extract the 90-percentile of wave speed from each recording (n=9 for myocyte monoculture, n=19 for co-cultures), comparing the two groups of numbers using two-sample Kolmogorov-Smirnov test. Our automated conduction velocity measurement system yields lower values than measurements of planar waves in paced tissue, which may be due to the inclusion of slowly moving highly curved wavefronts in the calculations (e.g. wave fronts near the core of a spiral wave). We note that the maximal observed wave velocity (measured manually) in our preparations was 185 mm/second, which is similar to planar wave conduction velocities in isotropic tissue reported by other groups.

### Simultaneous patch clamp electrophysiology-video recording

Current clamp was used to record electrophysiological traces from neurons, while video recording of cardiac behaviour was performed simultaneously using a CCD camera. The signal was recorded and analysed using custom-made procedures in Igor Pro (Wavemetrics). Image series were analysed after processing the images so that the value of each pixel p at frame t, denoted p(t), is replaced by the absolute value of p(t) − p(t−5).

Current clamp recordings were performed with a Multiclamp 700B amplifier (Molecular Devices). Borosilicate glass pipettes were filled with an internal solution containing (in mM): 110 potassium-gluconate, 40 HEPES, 2 ATP-Mg, 0.3 GTP and 4 NaCl (pH 7.2–7.3; osmolarity 270–285 mOsmol). Standard artificial cerebrospinal fluid (ACSF) bubbled with 5% CO2 was used as the bath solution, containing (in mM): 126 NaCl, 3 KCl, 1.25 NaH2PO4, 2 MgSO4, 2 CaCl2, 26 NaHCO3, and 10 glucose, pH 7.2–7.4. All recordings were conducted at 37°C. Square pulses of current or current steps were administered to depolarise the neurons and elicit action potential firing. Data were low-pass filtered at 2 kHz and acquired at 5 kHz.

### Proteomics

#### (i) In-gel trypsin digestion

Cell lysates were mixed with an equal volume of reducing SDS-PAGE sample buffer, heat denatured (95°C for 10 min) and run into the upper part of a Tris-Glycine 8-16% gel (185V, 10m min). Protein material was excised after Coomassie blue staining and cut into 1 – 2 mm^3^ gel pieces, which were placed into 1.5 mL sample tubes. Gel pieces were rinsed twice with wash solution for 18h in total (200 μl, 50% methanol, 5% acetic acid). The solutions were removed and gel pieces were dehydrated in acetonitrile (200 μl, 5 min). Supernatant were removed and gel pieces were dried in a vacuum centrifuge for 3 min. Disulfide reduction was performed with 10 mM DTT (30 μl) for 0.5 h, followed by alkylation with 100 mM iodoacetamide (30 μl) for 0.5 h. Supernatants were removed from the gel samples and dehydration with acetonitrile and evaporation performed as described above. Gel pieces were washed with 100 mM ammonium bicarbonate (200 μl, 10 min). Supernatants were removed and dehydration performed with acetonitrile and evaporation as above. The gel samples were then rehydrated on ice with freshly prepared trypsin solution (30 μl, 20 ng/μl sequencing grade trypsin [Promega] in 50 mM ammonium bicarbonate). After rehydration excess trypsin solution was removed and 50 mM ammonium bicarbonate (10 μl) was added to prevent dehydration of gel pieces. Gel samples were digested at 37°C for 18h. The gel pieces were then extracted sequentially with 50mM ammonium bicarbonate (60 μl), 50% acetonitrile, 5% formic acid (60 μl) and 85% acetonitrile, 5% formic acid (60 μl). The combined extracts were evaporated in a vacuum centrifuge and were redissolved in 5% acetonitrile, 0.1% formic acid (20 μl) on an ultrasonic bath and transferred into LC-MS sample vials.

#### (ii) Proteomics analysis by liquid chromatography-tandem mass spectrometry (LC-MS/MS)

Trypsin digested samples were analyzed either on an Amazon Ion Trap mass spectrometer (Bruker Daltonics) as described previously [47], or on Q-Exactive (Thermo Scientific) Hybrid Quadrupole-Orbitrap mass spectrometer LC-MS/MS system as detailed below. Liquid chromatography: A Dionex UltiMate3000 RSLCnano pump at 300 nL/min flow rate was used for analytical separation of digested peptides. A C18, 75 µm x 50 cm (2.6 µm particle size, 150 A; part number: 16126-507569 Thermo Scientific) analytical column was used for separation of peptides and the column temperature was 50°C. Mobile phase used was: A – 0.1% v/v formic acid in water (LC-MS grade) and B – 0.1% v/v formic acid in acetonitrile (LC-MS grade). Two hours linear gradient was used as below, a multistep gradients between 121-132 minutes were used to remove any carryover.

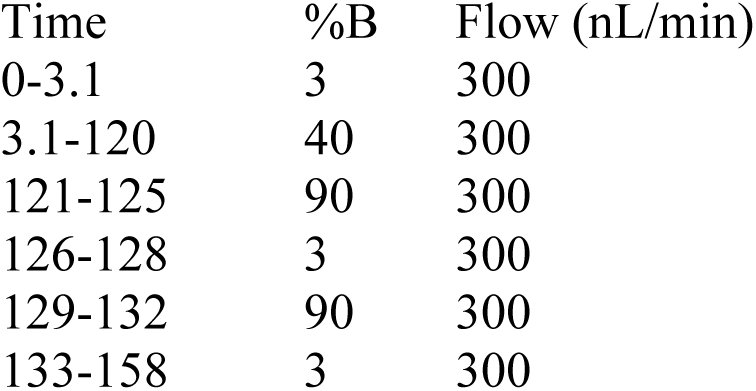

| Time | %B | Flow (nL/min) |
| --- | --- | --- |
| 0-3.1 | 3 | 300 |
| 3.1-120 | 40 | 300 |
| 121-125 | 90 | 300 |
| 126-128 | 3 | 300 |
| 129-132 | 90 | 300 |
| 133-158 | 3 | 300 |

A Dionex UltiMate 300 RS pump was used for desalting the peptide. A C18 PepMap (µ-Precolumn, 300 µM I.D. x 5 mm, 100 µm particle size, 100 A; Part number: 160454, Thermo Scientific) trapping column was used for trapping peptides. Buffer used for trapping and desalting peptides was 0.05% v/v Trifluoroacetic acid (TFA) in water (LC-MS grade). 1 µl of samples was injected and allowed for desalting for 3 minutes at 10 µl/min flow rate. After 3 minutes of desalting the trapping valve was changed and directed (backward flush) to analytical column for separation of all trapped peptides on trapping column.

#### (iii) Mass spectrometry

A bench top Q-Exactive (Thermo Scientific) Hybrid Quadrupole-Orbitrap mass spectrometer was used for data acquisition. A Top10 data dependent acquisition (DDA) method was used. Mass spectrometer was calibrated, for mass accuracy using positive ion calibration mixture provided by mass spectrometer manufacturer, prior to data acquisition. The conditions for the DDA mode were; Chromatographic peak width: 10 s, the Full MS conditions used-resolution: 70,000, AGC target: 1e6, maximum IT (injection time): 100 ms, scan range: 300 to 2000 m/z. The dd-MS2 conditions - resolution: 17,500. The AGC target conditions-5e4, maximum IT: 100 ms, loop count: 10 (i.e. Top 10), isolation width: 1.6 m/z, fixed first mass: 120.0 m/z, and the data dependent (dd) settings -under fill ratio: 10% (it sets up a minimum intensity threshold of 5e4 ions), charge exclusion: unassigned, 1, 8, >8, peptide match; preferred, dynamic exclusion: 30 s. Normalised Collison Energy (NCE) of 27 was used for fragmentation of peptides in a high-energy collision dissociation (HCD) cell. This method allows the selection, fragmentation and detection of ten precursors in a duty cycle time of 1.42 s.

#### (iv) Data processing and database searching

For the analyses carried out using the Amazon Ion Trap (Bruker Daltonics), raw LC-MS/MS data were processed and Mascot compatible files were created using DataAnalysis 4.0 software (Bruker Daltonics). Database searches were performed using the Mascot algorithm (version 2.5.1) and the UniProt_SwissProt database with taxonomy restriction ‘rat’ (v2015.02.04, number of entries 547,357, after taxonomy filter: 7,930). The following parameters were applied: 2+, 3+ and 4+ ions, peptide mass tolerance 0.3 Da, 13C = 2, fragment mass tolerance 0.6 Da, number of missed cleavages: two, instrument type: ESI-TRAP, fixed modifications: Carbamidomethylation (Cys), variable modifications: Oxidation (Met). For the analyses carried out using the Q-Exactive (Thermo Scientific) Hybrid Quadrupole-Orbitrap mass spectrometer, LC-MS/MS data (.raw files) were converted to .mgf files and Database searches were performed using the Mascot algorithm (version 2.5.1) and the UniProt_SwissProt database with taxonomy restriction ‘rat’ (v2015.11.26, number of entries after taxonomy filter: 7,954). The following parameters were applied: 2+, 3+ and 4+ ions, peptide mass tolerance 10 ppm, 13C = 2, fragment mass tolerance 0.06 Da, number of missed cleavages: two, instrument type: Q-Exactive, fixed modifications: Carbamidomethylation (Cys), variable modifications: Oxidation (Met).

Protein ratios (co-cultures/myocytes) were calculated from label-free intensities for all identified protein hits (see Fig S6). We chose a 3-fold change as a threshold for regulation.

### Insights from Proteomics

Proteomics highlights changes in cytoskeletal, metabolic and nuclear proteins which occur in co-cultures: To further shed light on the molecular mechanisms governing the difference in macroscopic behaviour, we looked at the protein expression profiles between the two cultures. We performed a comparative proteome analysis of harvested cell cultures. Database searches of merged peak list files were carried out using the MASCOT algorithm with taxonomy restriction to rat sequences. False-discovery rate (FDR) estimation was performed through a target-decoy search strategy and all results were displayed at a FDR of ≤1% (Supplement Table 1).

The identified proteins included many abundant cytoskeletal proteins (actin, tubulin, vimentin, vinculin, actinin), metabolic enzymes (alpha-enolase, GAPDH, pyruvate kinase, ATP synthase) and nuclear proteins (histones H2A, H2B, H3, H4, prelamin) which were detected commonly across the two samples.

Among the most strongly regulated protein hits we see a significant increase in several biological processes (See Fig S6 (A) and (B) from 2 independent experiments). From the STRING [48] and Expressence analysis, the most abundant class of proteins in the co-cultures can be seen in Supplement Table 2, Enrichment Analysis for two independent samples (SD5 and SD22, significance <0.05). Interestingly, using the Cellular Component analysis (Supplement Table 2, tab abbreviated CC), we observe a significant up regulation in translational and mitochondrial related proteins including mitochondrial respiratory chain complex III (Supplement Table 2, tab CC, and visualised in Figure S1 (C) and (D)).

### Opto-chemical and Opto-genetic neuronal stimulation and the effect of neuro transmitter release on cardiomyocytes

#### Caged-Nicotine

Electrophysiological experiments studying the effects of noradrenaline release by sympathetic neurons on cardiac behaviour employ methods that flow on drug and monitor voltage or calcium changes either via the patch pipette or beat rate changes using video microscopy. We first tested the co-cultures using conventional pharmacological approaches (with Sigma 50 µM nicotine stimulation), to demonstrate the expected results of noradrenaline release by stellate sympathetic cardiac neurons on myocytes. Conventional methods such as the aforementioned do not offer any temporal or spatial control. Such a technique does not bode well with high-throughput imaging methods as the drug effect and the imaging timings cannot be precisely coordinated.

Due to temporal limitations of drug administration and effects, an alternative technique suitable to high-throughput measurements employing caged compounds was employed. Caged compounds are light-sensitive probes that functionally encapsulate molecules first introduced in the 1970’s. These molecules are in an inactive form and upon irradiation, liberate the trapped molecule allowing targeted perturbation of physiological processes [49]. Caged nicotine known as RuBiNic (RuBi-Nicotine, Cat No. 3855, Tocris) has been used in neurotransmission studies [43]. Uncaging can be extremely rapid, controlled in time or space and quantitatively controlled and repeated. We tested the following neuron-myocyte densities: 1:5, 1:20 and 1:100,000 (see Figure 4 J for immuno-histochemistry staining of neurons in co-culture).

#### Neuronal optogenetics actuation

##### Optogenetic modification of neurons

Adenoviral vectors containing the transgene for hChR2(H134R)-eYFP were prepared in collaboration with the Stony Brook University Stem Cell Centre based on the expression cassette of the plasmid pcDNA3.1/hChR2(H134R)-eYFP (#20940; Addgene)[50]. Neurons were infected using Ambrosi’s method [50] with an optimised dose of adenovirus (multiplicity of infection (MOI) 2000) at 37°C for 2 h. Neuron expressing ChR2 was confirmed by eYFP reporter visualization. Following infection, fresh culture medium was added with a medium change at 12 h and then every 48 h with functional measurements performed on the sample from day 5 onwards. Optical recording of membrane voltage, Vm, was performed using the synthetic voltage-sensitive dye Di-4-ANBDQBS, spectrally compatible with ChR2. For Optogenetic experiments: Optical stimulation (470 nm) was provided at pulse lengths of 5–20 ms, at 0.5–8 Hz, using irradiances of 0.4–7 mW mm−2, as needed.

##### (i) Beat rate changes in response to neuronal ChR2 stimulation

Figure 4 illustrates OptoDyCE imaging (B) and the protocols of “optoelectrical” (via ChR2) vs “optochemical” (via Caged Nicotine) neural stimulation. Rat cardiac sympathetic stellate neurons are optogenetically transformed to respond to light activation, this promotes action potential firing in myocytes (Fig 4 D). Figure 4 (J i-iv), ChR2 expression by eYFP reporter (green) at four different neuronal plating densities 1:5, 1:20, 1:100 and 1:100,000. Optogenetic stimulation via ChR2: (Figure S5C) in the myocyte only culture shows no beat rate response, where as in the co-culture (Figure S5D), we observe a beat rate response during the periods of light stimulation. Figure S5(E), post processed traces using using custom-written Matlab software, where green is the raw trace, magenta is the trace after baseline subtraction and de-baselined green, blue is the magenta trace after median filtering, red indicates detected spike times, black is an indicator of when light is present (black down=light on).

Optochemical stimulation of neurons with nicotine (Figure S6Bii: pre-uncaging, Biii: post-uncaging, Biv: returning to baseline after several minutes of uncaging) and optoelectrical stimulation of neurons via ChR2 (Figure S5E), demonstrate functional coupling between neurons and myocytes in co-cultures. We additionally test this coupling by traditional pharmacological antagonists (nicotine, Figure S7). Genetically modified sympathetic neurons with ChR2 co-cultured with myocytes can be optically stimulated to drive a monolayer of myocytes (Figure S5Fi) and this effect can be blocked using a beta-blocker (metoprolol, Sigma Aldrich, Figure S5Fii).

##### (ii) Cardiomyocyte response to ChR2 stimulation with ChR2-expressing neurons as a function of neuron density

Stimulating the ChR2-CSSN (cardiac sympathetic stellate neurons) elicits a rate response in the cardiac monolayer. The number of neurons innervating the myocytes affects the firing frequency of myocytes. Different neuronal plating densities influence the cardiac response. We observe a dose-dependent effect (i.e. the greater the number of neurons innervating the myocytes, the greater the effect; Figure 4 I).

Firing frequency versus concentration of neurons in co-cultures: There is a fairly strong dose-dependent effect observed, (Figure 4 I). We observe that most values are >0, indicating a speed up. Where x axis represents ratios of neurons to myocytes (0.2=1:5, 0.05=1:20, 0.01=1:100, 0.001=1:10,000, 1e-05=1:100,000 and 0= myocytes/controls). No significant increase/decrease observed with light stimulation in controls (no neurons) versus co-cultures. We tested for reproducibility of light stimulation and precision (Fig 4 L). All data analysed using custom-written Matlab script.

##### (iii) Validation of a co-culture model and translational value-iPSc-CM and iPSc-peripheral neurons

The data we present are more anecdotal due to the limited n numbers. Example traces, and response to caged nicotine and optogenetic manipulation of iPSc-derived peripheral neurons with immunostains are presented to demonstrate the utility of the methodology presented in this paper. Cardiomyocyte response to ChR2 stimulation with ChR2-expressing peripheral neurons (MOI 2000) as a function of neuron density and blue light protocol of 3 sec on/off protocols were employed. Our data demonstrate the utility and power of our co-culture model and the applicability of OptoDyCE [22] for contactless opto-chemical and optogenetics experiments required in cell specific perturbations.

Cor.4U® human iPS-derived cardiomyocytes and Peri.4U® human iPS-derived peripheral neurons were grown as described by Axiogenesis handling guides (https://ncardia.com/resources/#manuals).

Here we performed direct adenoviral gene delivery in human iPSc-derived peripheral neurons (ChR2-hiPSc-PN, obtained from Axiogenesis) and observe the cardiac response to light stimulation of neurons (Figure S7), confirming that optogenetic stimulation of co-cultures is a suitable method and alternative for dissecting neural mediated cardiac function in a ‘petri-dish’. To confirm the myocyte-like properties of the iPS-dervied cardiomyocytes, we utilised mouse anti-α-actinin primary antibody and imaged them using a confocal microscope, using the Olympus FluoView FV1000 system. All samples were fixed in 3.7% formaldehyde after performing functional experiments. Cells were permeabilised using 0.02% TritonX-100 for 5 minutes. Antibodies were diluted using 1% bovine serum albumin (Amersham PLC, Amersham, UK). 1% FBS was used as a blocking agent. After antibody staining, cell nuclei were stained with 1 μg/mL DAPI with 10 min incubation in PBS. From gene chip data provided by Ncardia, the iPSc-peripheral neurons Peri.4U have a high expression of Th gene (7719 where 501-1000 = high expression levels); and very low expression levels of ChAT gene (123, where <150 = absent/very low expression).

## SUPPLEMENTARY FIGURES

**Figure S1:**
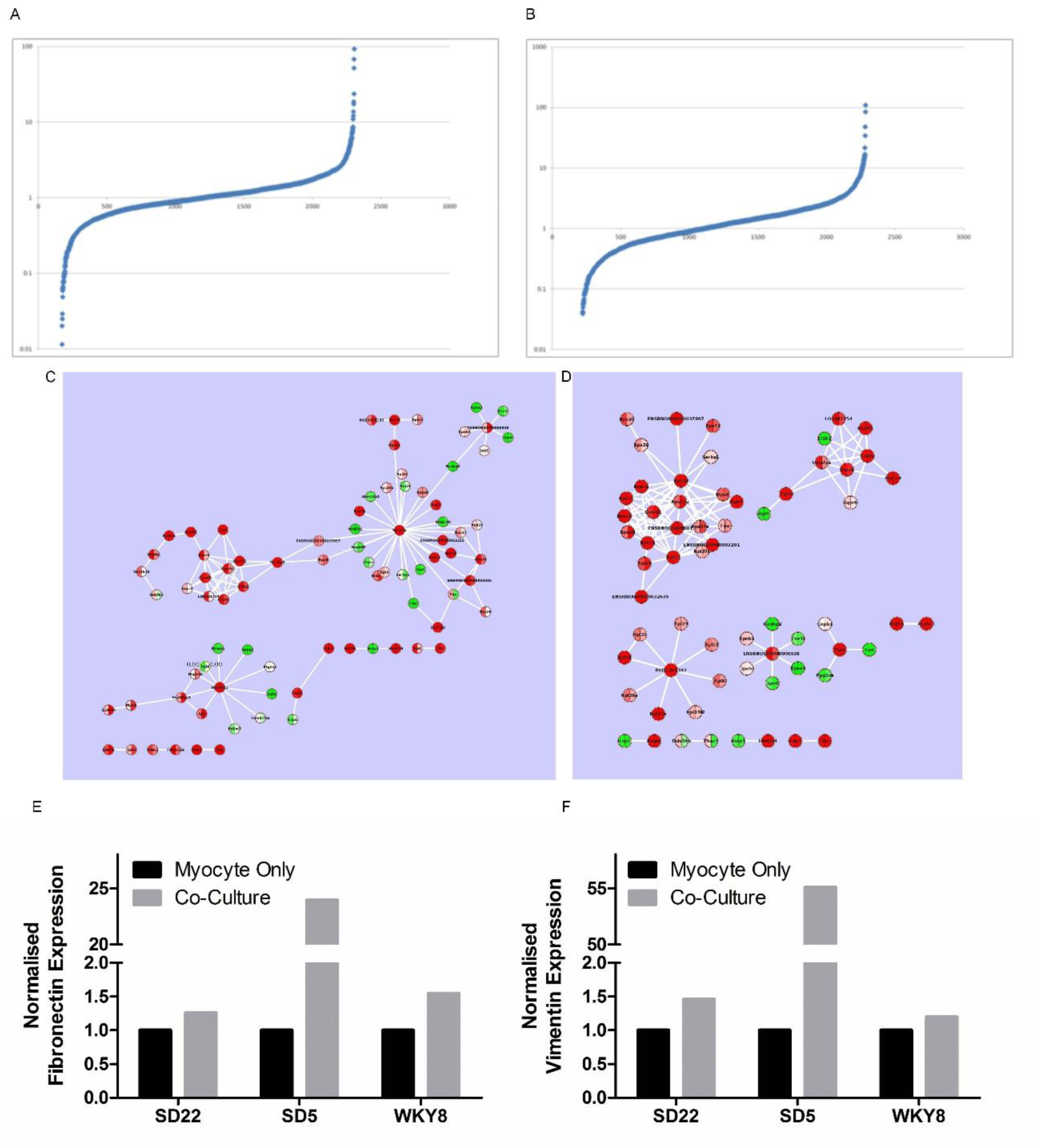
Quantitative, label-free proteomics and Western blot analysis on co-cultures. (A-B): In two independent experiments (SD22 and SD5) and their representative protein regulation levels, where individual peptides are identified by database search and quantified using label-free quantification. (C-D) Network visualization of co-cultures from the quantified proteins using Cytoscape (with text mining”/”no evidence” removed). Raw protein association data was obtained from STRING. Green nodes represent proteins which are at least 2 fold down-regulated (<0.5), red nodes are proteins which are at least 2 fold up-regulated (>2) and grey nodes are proteins which show less than two-fold variation (between 0.5 to 2). Results for the network condensation analysis using Expressence in SD22 and SD5 experiments independently. The modules in the figure represent the groups of highest correlated change for the larger network. The node colour is mapped to Measurement values (log transformed MaxQuant intensity values). (E-F) Western blot analysis and confirmation of protein regulation in two randomly selected proteins from independent experiments (SD22, SD5 and WKY8).

**Figure S2:**
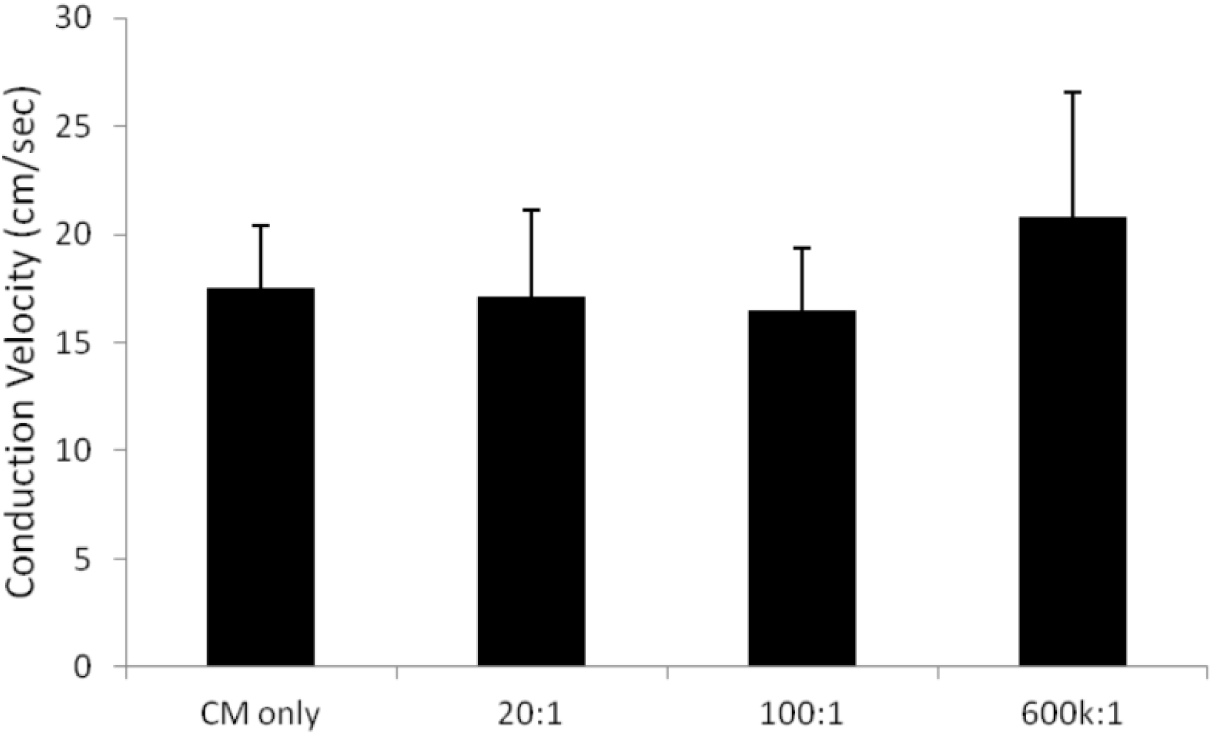
Effects of sympathetic cardiac stellate neurons on conduction velocity in well-connected cardiac cultures using optical mapping (Rhod-4 AM dye) and 1 Hz electrical stimulation. Overall, co-culturing myocytes with sympathetic neurons at 3 different concentrations of myocyte-neuron co-cultures did not alter conduction velocity of cardiomyocytes, where average conduction velocities (±std dev) were 17.505 ±2.92 in myocyte only cultures, 17.092 ±4.01 in 20:1 co-cultures, 16.46 ±2.91 in 100:1 co-cultures and 20.791 ±5.79 in 600k:1 co-cultures; (myocyte only cultures n=6, 20:1 co-cultures n=5, 100:1 co-cultures n=4, 600k:1 co-cultures n=6). Plot of conduction velocity of different co-cultures and myocyte only cultures.

**Figure S3:**
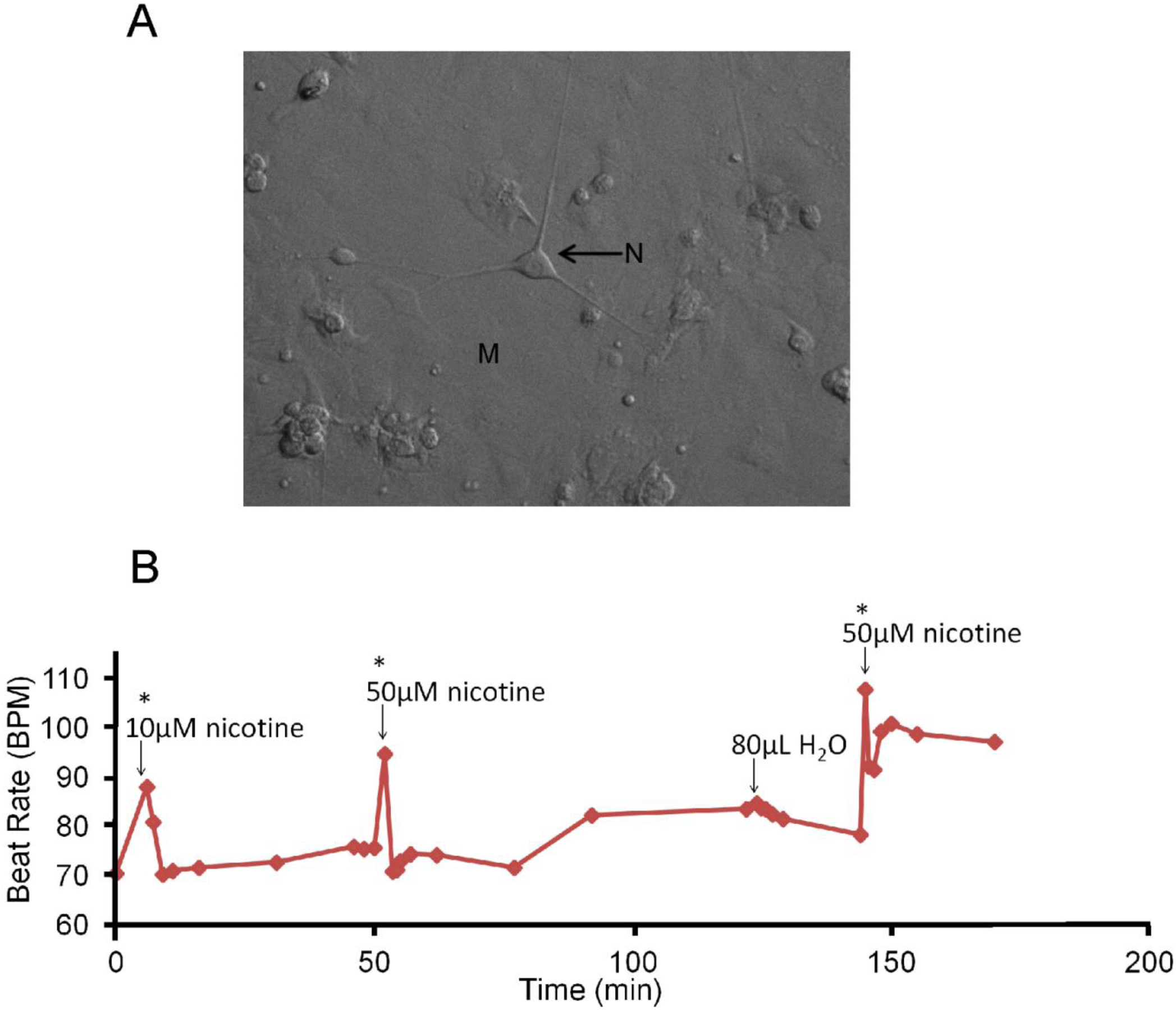
Response to nicotine (A) Bright field image of a confluent cardiac monolayer with cardiac sympathetic stellate neurons seeded on top. (B) An example trace of a co-culture nicotine stimulation experiment. Repeat nicotine doses (10 and 50 µM) caused transient increases in myocyte beat-rate, control vehicle of the same volume had no effect.

**Figure S4:**
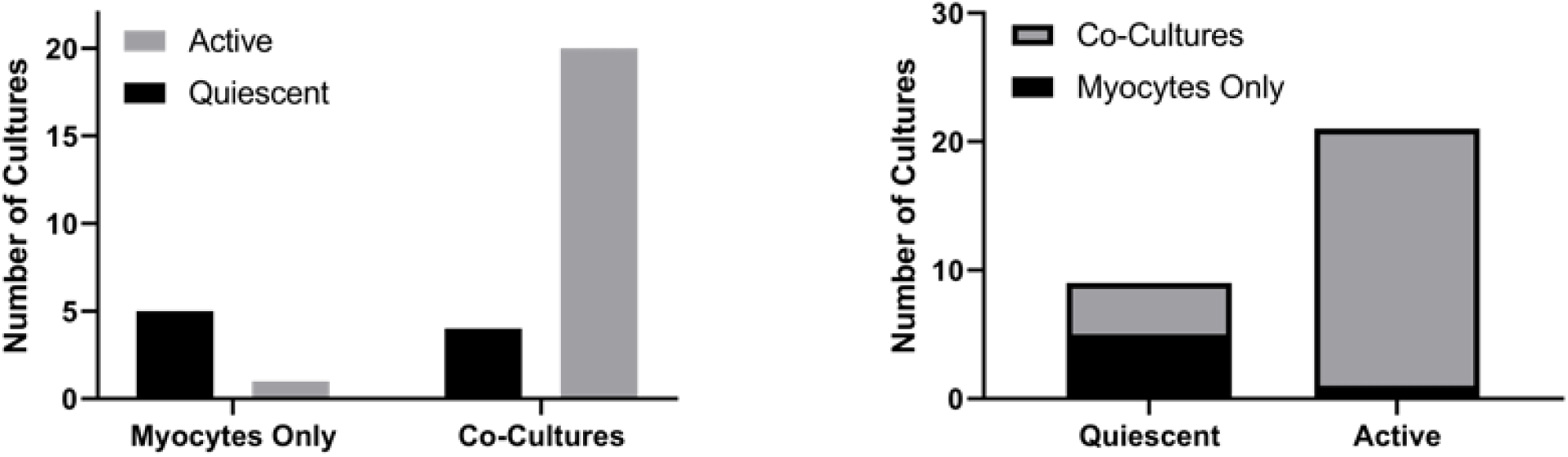
OptoDyce high throughput co-culture experiments to study the effects of sympathetic cardiac stellate neurons on cardiac activity in well-connected quiescent cardiac cultures using optical mapping (cultures loaded with dye Di-4-ANBDQBS). Fishers exact test (two sided), p= 0.0046 statistically significant. n=6 myocytes and n=24 for co-cultures.

**Figure S5:**
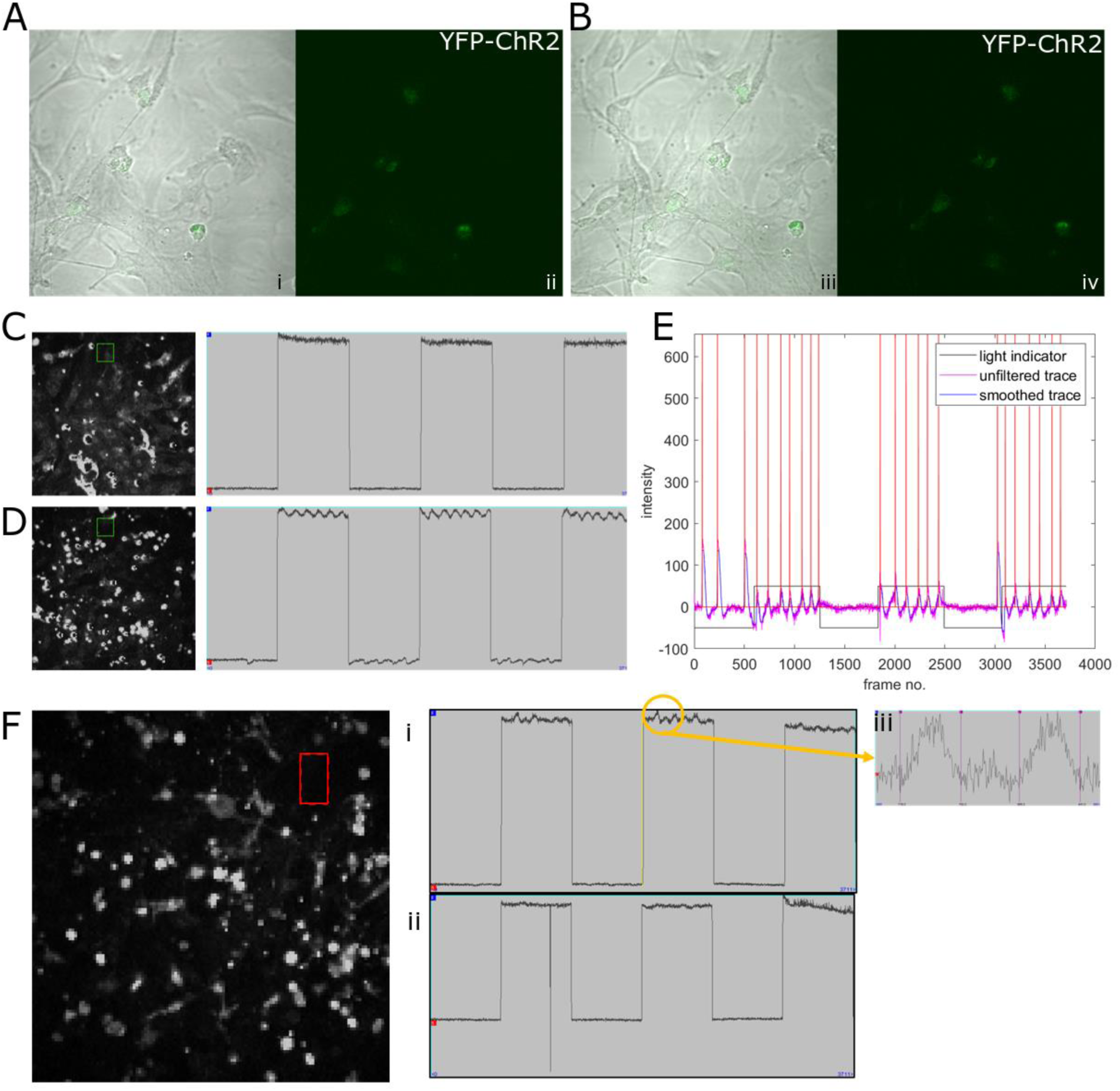
OptoDyce high throughput co-culture experiments to drive ChR2 stellate sympathetic neurons. Co-cultures of neurons and myocytes (loaded with dye Di-4-ANBDQBS spectrally compatible with ChR2). (A and B) Fixed bright field image and YFP-ChR2 (in the neurons) confocal image of a co-culture (1:5). (C-D) Neuronal stimulation via ChR2, top panel (C) myocyte only culture therefore no beat rate response observed to ChR2 stimulation, bottom panel (D), beat rate response observed during the periods of light stimulation in a co-culture. (E) Post processed traces using custom-written Matlab script. Magenta is the raw trace, blue is the trace after baseline subtraction and after median filtering, red indicates detected spike times, black is an indicator of when light is present (black down=light off). (F) Neural stimulation via ChR2. (i) Raw traces showing beat rate responses observed in co-cultures, and (ii) no beat rate responses observed in co-cultures that were blocked by the beta blocker metoprolol (10 μM). (iii) Zoomed in region from an action potential response.

**Figure S6:**
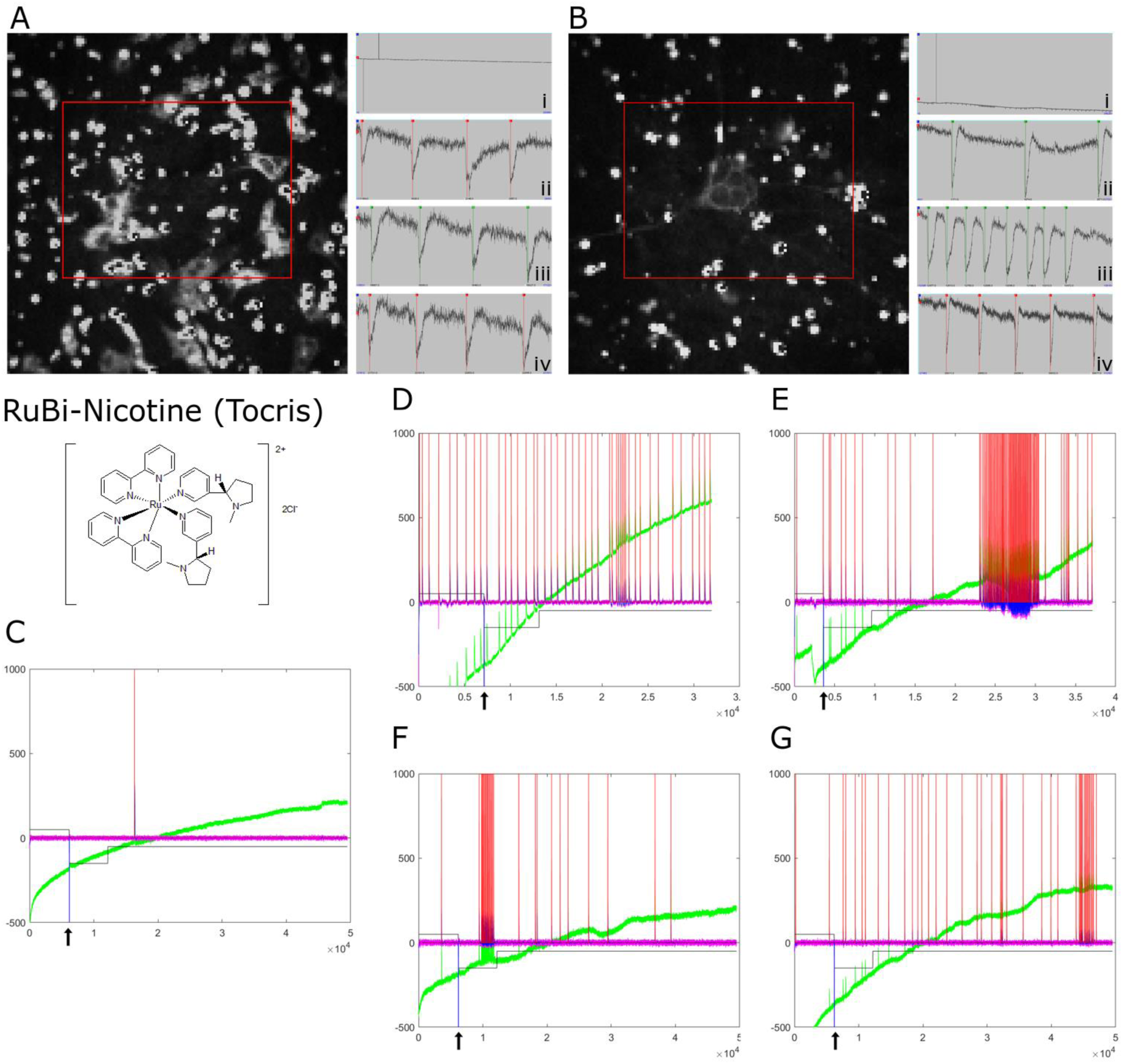
OptoDyCE high throughput cardiac imaging upon uncaging of nicotine (RuBi-Nicotine from Tocris). Panel (A) cardiac only monolayer (control, no neurons). Trace (Ai) full experimental trace, long spike corresponding to flash of light to uncage nicotine. (Bii) Control beat rate prior to uncaging (before light flash); (Biii) Post-nicotine uncaging beat rate (no change in beat rate); (Biv) Post-nicotine after 3 minutes (no change in beat rate). Panel (B) ROI of cluster of neurons sitting on a monolayer of cardiomyocytes. Trace (Bi), full experimental trace, long spike corresponding to flash of light to uncage nicotine. (Bii) Control beat rate prior to uncaging (before light flash); (Biii) Post-nicotine uncaging beat rate (speeds up and shows bursting behaviour); (Biv) Post-nicotine after 3 minutes. Example traces from the different uncaging experiments in the different neuron-myocyte concentration plates. (C) No responses to nicotine were detected in the control myocyte only dishes (representative example). (D) 1:5 neuron-mycoyte co-culture. (E) 1:20 neuron-myocyte co-culture. (F) 1:100 neuron-myocyte co-culture. (G) 1:100,000 neuron-myocyte co-culture. Arrow indicates moment of blue light flash to uncage the caged nicotine.

**Figure S7:**
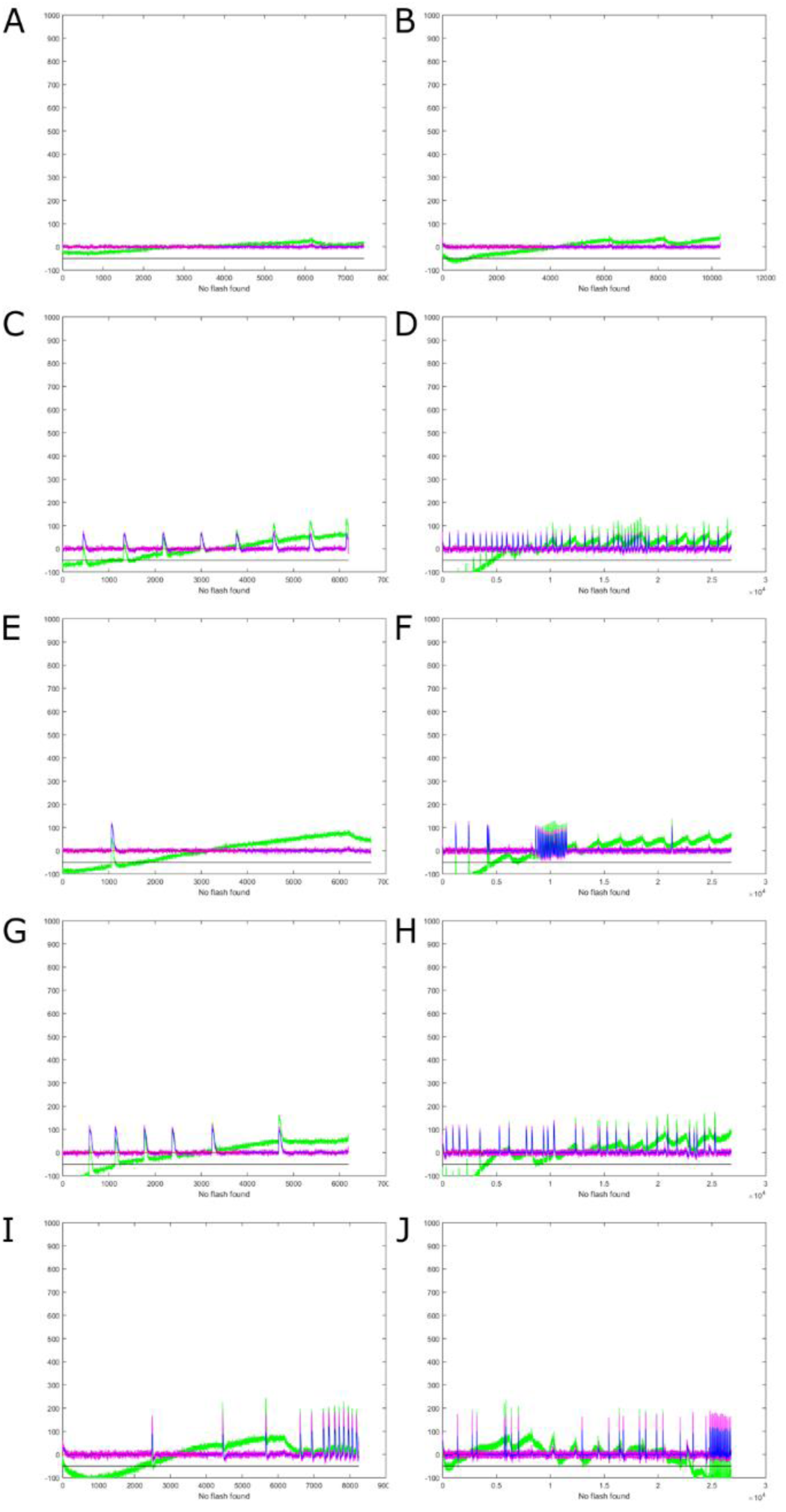
OptoDyce high throughput co-culture experiments to study the effect of standard nicotine on co-cultures. Co-cultures of neurons and myocytes at different neuron to myocyte ratios (A myocyte only, C 1:5, E 1:20, G 1:100 and I 1:100,000) were exposed to 10 µM nicotine (nicotine response traces B, D, F, H and J,) and the voltage traces were recorded (cultures loaded with dye Di-4-ANBDQBS). Example traces and responses are shown in this figure.

**Figure S8:**
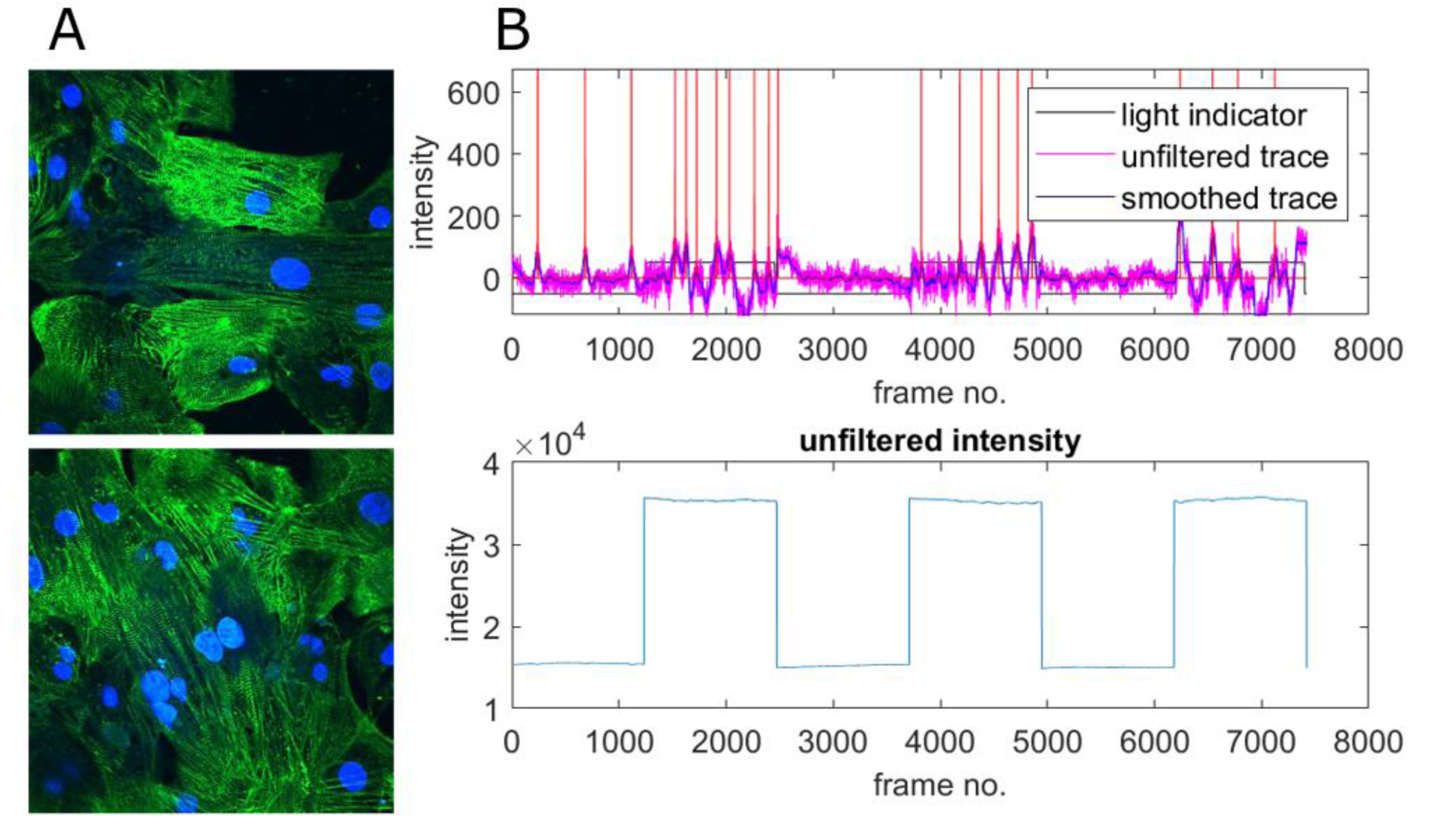
(A and B) Pilot co-cultures of Cor.4U hiPSC-derived cardiomyocytes and Peri.4U hiPSC-derived peripheral neurons (largely sympathetic) developed by Axiogenesis (now Ncardia). (A) Cardiomyocytes labelled with alpha-actinin staining (green) and DAPI nuclear stain (blue). (G) Example traces showing cardiac activity followed by long light pulse stimulation of the neurons, action potentials are evoked indirectly in the myocytes via the ChR2-light-sensitized neurons.

## References

1. Winfree, A.T., When time breaks down: The three-dimensional dynamics of electrochemical waves and cardiac arrhythmias. 1987, Princeton, N.J: Princeton University Press.

2. Bub, G. and R.A. Burton, Macro-micro imaging of cardiac-neural circuits in co-cultures from normal and diseased hearts. J Physiol, 2014. 2014: p. 14.

3. Bers, D.M., Calcium cycling and signaling in cardiac myocytes. Annu Rev Physiol, 2008. 70: p. 23–49.

4. Devinsky, O., Effects of Seizures on Autonomic and Cardiovascular Function. Epilepsy Curr, 2004. 4(2): p. 43–46.

5. Julius, S., Effect of sympathetic overactivity on cardiovascular prognosis in hypertension. Eur Heart J, 1998. 19(8): p. F14–8.

6. Cohn, J.N., et al., Plasma Norepinephrine as a Guide to Prognosis in Patients with Chronic Congestive Heart Failure. New England Journal of Medicine, 1984. 311(13): p. 819–823.

7. Chen, P.S., et al., Role of the autonomic nervous system in atrial fibrillation: pathophysiology and therapy. Circ Res, 2014. 114(9): p. 1500–15.

8. Chen, P.S., et al., Sympathetic nerve sprouting, electrical remodeling and the mechanisms of sudden cardiac death. Cardiovasc Res, 2001. 50(2): p. 409–16.

9. Cao, J.M., et al., Relationship between regional cardiac hyperinnervation and ventricular arrhythmia. Circulation, 2000. 101(16): p. 1960–9.

10. Boogers, M.J., et al., Cardiac sympathetic denervation assessed with 123-iodine metaiodobenzylguanidine imaging predicts ventricular arrhythmias in implantable cardioverterdefibrillator patients. J Am Coll Cardiol, 2010. 55(24): p. 2769–77.

11. Fallavollita, J.A., et al., Denervated Myocardium Is Preferentially Associated With Sudden Cardiac Arrest in Ischemic Cardiomyopathy: A Pilot Competing Risks Analysis of Cause-Specific Mortality. Circ Cardiovasc Imaging, 2017. 10(8).

12. Tomek, J., et al., β-Adrenergic receptor stimulation inhibits proarrhythmic alternans in postinfarction border zone cardiomyocytes: a computational analysis. American Journal of Physiology-Heart and Circulatory Physiology, 2017. 313(2): p. H338–H353.

13. Tomek, J., et al., β-Adrenergic Receptor Stimulation and Alternans in the Border Zone of a Healed Infarct: An ex vivo Study and Computational Investigation of Arrhythmogenesis. Frontiers in Physiology, 2019. 10(350).

14. Gardner, R.T., et al., Targeting protein tyrosine phosphatase sigma after myocardial infarction restores cardiac sympathetic innervation and prevents arrhythmias. Nat Commun, 2015. 6(6235).

15. Furshpan, E.J., et al., Chemical transmission between rat sympathetic neurons and cardiac myocytes developing in microcultures: evidence for cholinergic, adrenergic, and dual-function neurons. Proc Natl Acad Sci U S A, 1976. 73(11): p. 4225–9.

16. Horackova, M., et al., Cocultures of adult ventricular myocytes with stellate ganglia or intrinsic cardiac neurones from guinea pigs: spontaneous activity and pharmacological properties. Cardiovasc Res, 1993. 27(6): p. 1101–8.

17. Shaw, R.M. and Y. Rudy, Ionic mechanisms of propagation in cardiac tissue. Roles of the sodium and L-type calcium currents during reduced excitability and decreased gap junction coupling. Circ Res, 1997. 81(5): p. 727–41.

18. Kleber, A.G. and Y. Rudy, Basic mechanisms of cardiac impulse propagation and associated arrhythmias. Physiol Rev, 2004. 84(2): p. 431–88.

19. Tung, L. and Y. Zhang, Optical imaging of arrhythmias in tissue culture. Journal of Electrocardiology, 2006. 39(4): p. S2–S6.

20. Burton, R.A., et al., Optical control of excitation waves in cardiac tissue. Nat Photonics, 2015. 9(12): p. 813–816.

21. Shcherbakova, O.G., et al., Organization of β-adrenoceptor signaling compartments by sympathetic innervation of cardiac myocytes. The Journal of Cell Biology, 2007. 176(4): p. 521–533.

22. Klimas, A., et al., OptoDyCE as an automated system for high-throughput all-optical dynamic cardiac electrophysiology. Nat Commun, 2016. 7: p. 11542.

23. Tao, T., D.J. Paterson, and N.P. Smith, A model of cellular cardiac-neural coupling that captures the sympathetic control of sinoatrial node excitability in normotensive and hypertensive rats. Biophys J, 2011. 101(3): p. 594–602.

24. Himel, H.D.I.V., et al., Optical imaging of arrhythmias in the cardiomyocyte monolayer. Heart Rhythm, 2012. 9(12): p. 2077–2082.

25. Ogawa, S., et al., Direct contact between sympathetic neurons and rat cardiac myocytes in vitro increases expression of functional calcium channels. J Clin Invest, 1992. 89(4): p. 1085–93.

26. Atkins, D.L., et al., Regulation of rat cardiac myocyte growth by a neuronal factor secreted by PC12 cells. Pediatr Res, 1997. 41(6): p. 832–41.

27. Takeuchi, A., et al., Device for co-culture of sympathetic neurons and cardiomyocytes using microfabrication. Lab Chip, 2011. 11(13): p. 2268–75.

28. Larsen, H.E., K. Lefkimmiatis, and D.J. Paterson, Sympathetic neurons are a powerful driver of myocyte function in cardiovascular disease. Sci Rep, 2016. 6: p. 38898.

29. Claycomb, W.C., Biochemical aspects of cardiac muscle differentiation. Possible control of deoxyribonucleic acid synthesis and cell differentiation by adrenergic innervation and cyclic adenosine 3’:5’-monophosphate. J Biol Chem, 1976. 251(19): p. 6082–9.

30. Kreipke, R.E. and S.J. Birren, Innervating sympathetic neurons regulate heart size and the timing of cardiomyocyte cell cycle withdrawal. J Physiol, 2015. 593(23): p. 5057–73.

31. Oh, Y., et al., Functional Coupling with Cardiac Muscle Promotes Maturation of hPSC-Derived Sympathetic Neurons. Cell Stem Cell, 2016. 19(1): p. 95–106.

32. Coppen, S.R., et al., Comparison of connexin expression patterns in the developing mouse heart and human foetal heart. Mol Cell Biochem, 2003. 242(1-2): p. 121–7.

33. Giovannone, S., B.F. Remo, and G.I. Fishman, Channeling diversity: gap junction expression in the heart. Heart rhythm, 2012. 9(7): p. 1159–1162.

34. Luke, R.A. and J.E. Saffitz, Remodeling of ventricular conduction pathways in healed canine infarct border zones. J Clin Invest, 1991. 87(5): p. 1594–602.

35. Vaseghi, M. and K. Shivkumar, The role of the autonomic nervous system in sudden cardiac death. Prog Cardiovasc Dis, 2008. 50(6): p. 404–19.

36. Shen, M.J. and D.P. Zipes, Role of the autonomic nervous system in modulating cardiac arrhythmias. Circ Res, 2014. 114(6): p. 1004–21.

37. Bowes, J., et al., Reducing safety-related drug attrition: the use of in vitro pharmacological profiling. Nat Rev Drug Discov, 2012. 11(12): p. 909–22.

38. He, Y. and P.W. Baas, Growing and working with peripheral neurons. Methods Cell Biol, 2003. 71: p. 17–35.

39. Li, D., et al., Abnormal intracellular calcium homeostasis in sympathetic neurons from young prehypertensive rats. Hypertension, 2012. 59(3): p. 642–9.

40. Filevich, O., M. Salierno, and R. Etchenique, A caged nicotine with nanosecond range kinetics and visible light sensitivity. Journal of Inorganic Biochemistry, 2010. 104(12): p. 1248–1251.

41. Macgregor, A., et al., NAADP controls cross-talk between distinct Ca2+ stores in the heart. J Biol Chem, 2007. 282(20): p. 15302–11.

42. Hwang, S.-m., T.Y. Kim, and K.J. Lee, Complex-periodic spiral waves in confluent cardiac cell cultures induced by localized inhomogeneities. Proceedings of the National Academy of Sciences of the United States of America, 2005. 102(29): p. 10363–10368.

43. Filevich, O., M. Salierno, and R. Etchenique, A caged nicotine with nanosecond range kinetics and visible light sensitivity. J Inorg Biochem, 2010. 104(12): p. 1248–51.

44. Ambrosi, C.M., et al., Optogenetics-enabled assessment of viral gene and cell therapy for restoration of cardiac excitability. Sci Rep, 2015. 5: p. 17350.

45. Jia, Z., et al., Stimulating cardiac muscle by light: cardiac optogenetics by cell delivery. Circ Arrhythm Electrophysiol, 2011. 4(5): p. 753–60.

46. Bozzola, J.J., Conventional specimen preparation techniques for scanning electron microscopy of biological specimens. Methods Mol Biol, 2007. 369: p. 449–66.

47. Kar, P., et al., Dynamic assembly of a membrane signaling complex enables selective activation of NFAT by Orai1. Current biology : CB, 2014. 24(12): p. 1361–1368.

48. Jensen, L.J., et al., STRING 8--a global view on proteins and their functional interactions in 630 organisms. Nucleic Acids Res, 2009. 37(Database issue): p. D412–6.

49. Ellis-Davies, G.C., Caged compounds: photorelease technology for control of cellular chemistry and physiology. Nat Methods, 2007. 4(8): p. 619–28.

50. Ambrosi, C.M. and E. Entcheva, Optogenetic control of cardiomyocytes via viral delivery. Methods Mol Biol, 2014. 1181: p. 215–28.

