## Supplemental Video-3 for "Optical interrogation of sympathetic neuronal effects on macroscopic cardiac monolayer dynamics"

### Slide 1
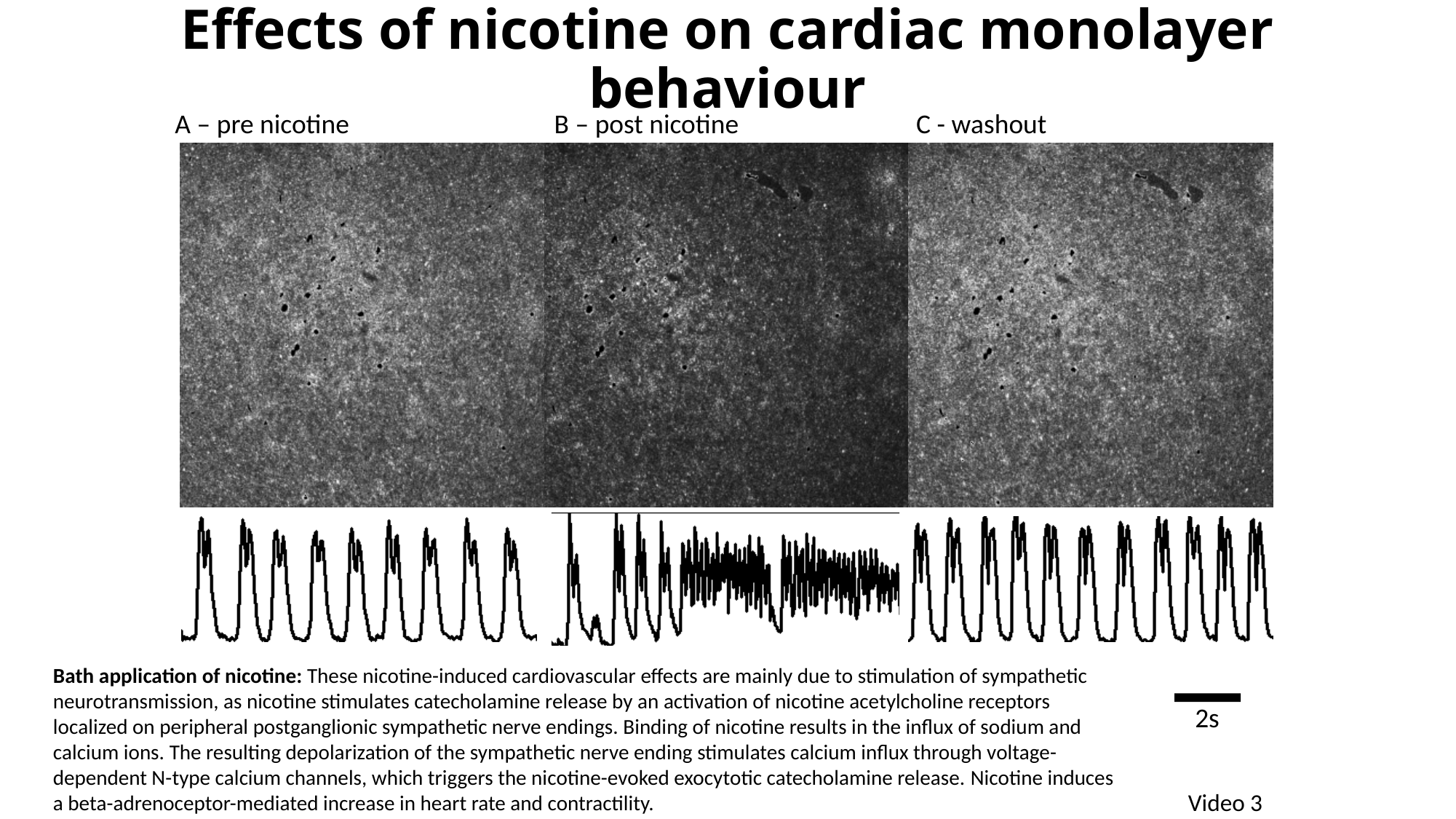

#### Effects of nicotine on cardiac monolayer behaviour
A – pre nicotine
B – post nicotine
C - washout
Bath application of nicotine: These nicotine-induced cardiovascular effects are mainly due to stimulation of sympathetic neurotransmission, as nicotine stimulates catecholamine release by an activation of nicotine acetylcholine receptors localized on peripheral postganglionic sympathetic nerve endings. Binding of nicotine results in the influx of sodium and calcium ions. The resulting depolarization of the sympathetic nerve ending stimulates calcium influx through voltage-dependent N-type calcium channels, which triggers the nicotine-evoked exocytotic catecholamine release. Nicotine induces a beta-adrenoceptor-mediated increase in heart rate and contractility.
2s
Video 3
